# Stemness factor Mex3a times translation and protein trafficking to ensure robust differentiation of olfactory sensory neurons

**DOI:** 10.64898/2025.12.26.696633

**Authors:** Martín Escamilla del Arenal, Lauren C. Tang, Albana Kodra, Hani Shayya, Aileen Ugurbil, Olga Stathi, Keskin Abdurrahman, Adan Horta, Joan Pulupa, Junqiang Ye, Marko Jovanovic, Stavros Lomvardas, Rachel Duffié

## Abstract

During the switch from progenitor to differentiated cell, cellular physiology must change to accommodate increased translation and trafficking of membrane-bound proteins. We identify RNA-binding and E3 ubiquitin ligase Mex3a as a key driver of proper neuronal differentiation by regulating mRNA translation and trafficking of cell surface proteins in the context of Unfolded Protein Response (UPR) signaling. Loss of Mex3a in immature olfactory sensory neurons (OSNs) leads to defects in cilia structure, cell surface protein expression, and planar cell polarity in mature OSNs. Proteomics reveal a Mex3a-dependent decrease in proteins related to vesicle transport, lipid metabolism, and ribosome biogenesis. We identify RNA and ubiquitin targets of Mex3a and provide evidence that Mex3a may confer K27 ubiquitin linkage on substrates. Finally, modulating cellular levels of Mex3a changes the recruitment of translation factors Serbp1 and p-eEF2 to ribosomes with possible effects on translation. Our data reveal how a stemness factor regulates development post-transcriptionally and post-translationally to ensure robust differentiation.

**Highlights:** Loss of stemness factor Mex3a in immature olfactory neurons leads to defects in mature olfactory neurons.

Translation/Trafficking of cell surface proteins, cilia structure, and planar cell polarity are compromised in the absence of Mex3a.

Mex3a may confer K27 ubiquitination on stress granule protein Serbp1 and ribosome protein Rps7.

Mex3a levels are associated with Serbp1 and p-eEF2 recruitment to ribosomes.

## Introduction

Many terminally differentiated cell types interphase with the exterior world in order to sense, protect from, or consume the environment (such as sensory, immune, and intestinal cells). As cells differentiate, higher levels of “outward” facing, cell surface genes are expressed, in contrast to “inward” facing genes such as transcription factors or cell signaling genes, which are expressed in stem cells and constitutively^1^. During differentiation, then, the physiology of a cell must change dramatically to accommodate the translation and trafficking of these surface or secreted proteins. The endomembrane system must expand^2^, and cytoskeletal structures must form to transport cargo and support changes in cell shape. Neurons provide a clear example of this: neural progenitors give rise to highly specialized cells with axons and dendrites that emit and perceive chemical and electrical signals. The processes that regulate the developmental switch from “inward” facing stem cells to “outward” facing differentiated cells are still incompletely understood.

The evolutionarily conserved^3–5^ stemness factor Mex3a is expressed in stem cells of the intestinal crypt^6^ and is a hallmark of many cancers^7^. We recently reported Mex3a expression in the immature neurons of the main olfactory epithelium (MOE)^8^. Located in the nasal cavity, the sensory neurons of the olfactory epithelium regenerate throughout life^9,10^. Whereas Mex3a expression is downregulated in other brain regions due to age-related differentiation^5,11^, olfactory sensory neurons (OSNs) are an excellent model to study Mex3a function due to characteristic life-long neuro-regeneration. In our previous study, we found that Mex3a post-transcriptionally represses olfactory receptors. In addition, loss of Mex3a leads to higher levels of UPR transcription factor Atf5 and downstream targets. Intriguingly, despite Mex3a’s exquisite spatiotemporal restriction to immature neurons, we observe lasting effects on mature neurons upon loss of Mex3a, including axon guidance defects and skewed OR choice^8^.

Here we provide evidence that loss of Mex3a during OSN differentiation leads to defects in cilia structure, membrane-bound protein trafficking, and planar cell polarity in mature OSNs. Proteomics in mature OSNs reveal that Mex3a cKO neurons exhibit reduced levels of proteins related to lipid metabolism, ribosome biogenesis, and vesicle trafficking. We identify putative RNA and ubiquitin targets of Mex3a in the MOE, and we propose that Mex3a confers K27 ubiquitin chains on its targets, in support of a non-proteolytic role for Mex3a-mediated ubiquitination. Finally, we identify a link between Mex3a, translation factor eEF2, and translation associated factor Serbp1. We propose that they cooperate to fine-tune translation rates during and after the Unfolded Protein Response (UPR) signaling pathway is activated in OSNs.

## Results

### Loss of stemness factor Mex3a leads to deficits in cell surface protein trafficking and translation, cilia structure, and planar cell polarity in mature neurons

We previously identified Mex3a, a KH and RING domain containing protein, to be expressed specifically in immature neurons in the olfactory epithelium^8^. Immunofluorescence in wildtype (WT) mice reveals non-overlapping expression with mature olfactory sensory neuron marker and endoplasmic reticulum (ER) protein Calmegin^12^ (Figure 1A). Conditional knockout of Mex3a using a Mex3a flox mouse^13^ with the Foxg1iresCre^14^ driver (referred to as Mex3a cKO throughout) leads to loss of Mex3a protein in immature olfactory sensory neurons (Figure 1A). In the absence of Mex3a, immature neurons exhibit accelerated translation of olfactory receptor (OR) proteins^8^, the seven transmembrane G protein coupled receptors (GPCRs) that are critical to OSN identity and function. However, in mature OSNs differentiated from Mex3a cKO immature neurons, OR expression is significantly reduced at the protein level (Figure 1B-C), suggesting a deficit in mRNA translation, protein trafficking, or protein stability of this membrane-bound gene family. All four ORs tested by immunofluorescence exhibited decreased protein expression. On the other hand, at the mRNA level in mature OSNs, two of these ORs showed no significant change, one OR was significantly increased and one significantly decreased (data not shown). This significant decrease in OR protein level was specific to apical, mature neurons, and not to basal, immature neurons (Figure S1A-B). Curiously, we also observe reduced OR immunoreactivity in OSN cilia (Figure 1B, magenta arrows), supporting a defect in trafficking of OR proteins. To determine whether reduced expression level was specific to OR proteins, or if this phenotype extended to other proteins, we stained for another membrane-bound, cilia protein expressed in mature OSNs, Adcy3^15^. Adcy3 also exhibited significantly reduced fluorescence intensity in the Mex3a cKO compared to WT littermates (Figure S1C-E), raising the possibility that cilia-targeted and/or membrane-bound proteins exhibit reduced expression due to defects in mRNA translation protein trafficking, or protein stability in mature OSNs upon loss of Mex3a in immature OSNs.

**Figure 1:**
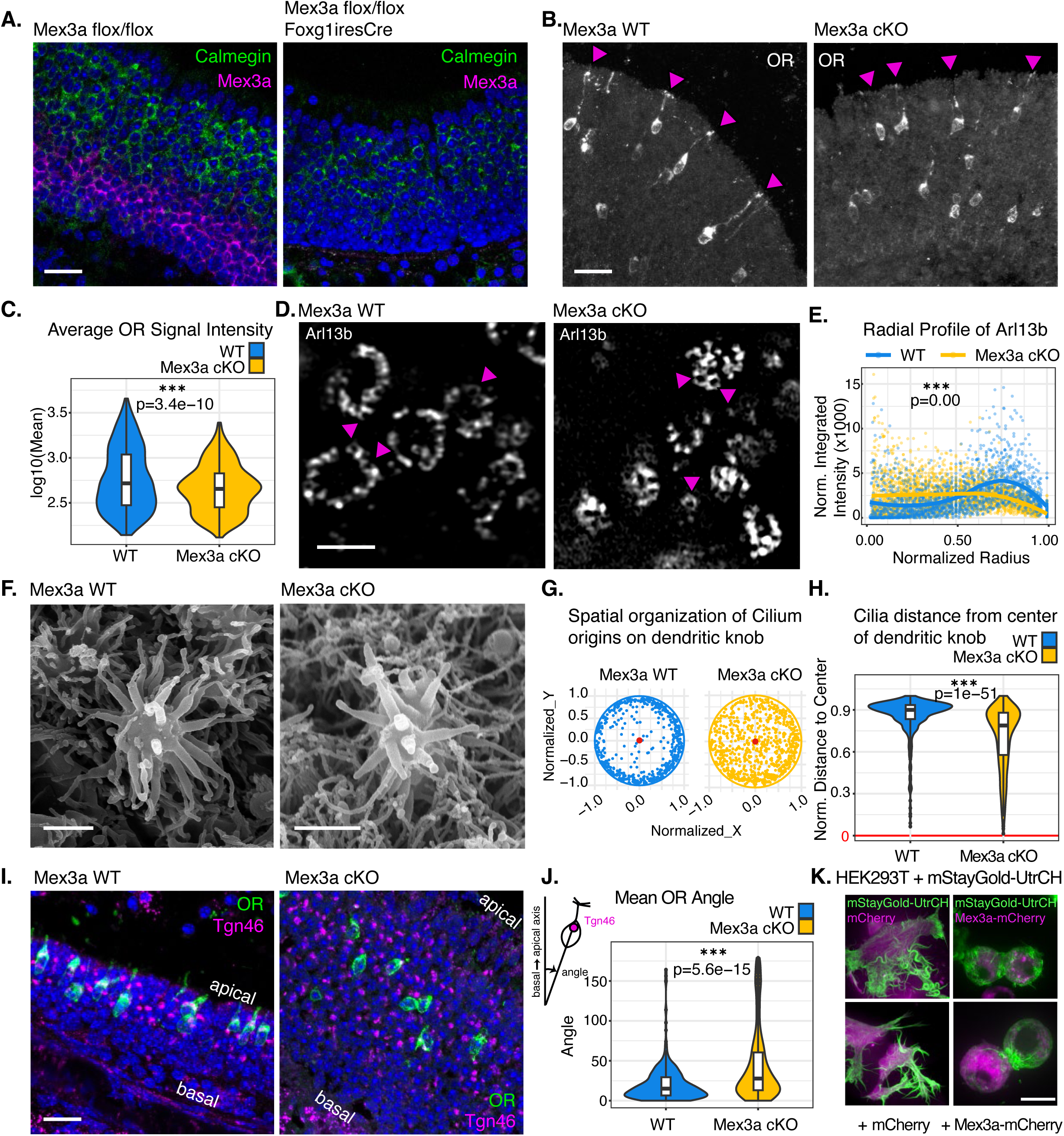
Loss of stemness factor Mex3a leads to deficits in cell surface protein trafficking and translation, cilia structure, and planar cell polarity in mature neurons. **A.** Immunofluorescence for Mex3a (magenta) and Calmegin (green) in Mex3a WT (left) and cKO (right) main olfactory epithelium, three-week-old littermates. Representative image from at least 5 biological replicates. DAPI in blue. Scale bar 25 µM. **B.** Representative immunofluorescence image for OR protein (pooled antibodies against C7, M71, and P2) in white. Staining in cilia highlighted with magenta triangles. Scale bar 25 µM. **C.** Violin plot quantifying OR cell body protein signal intensity per OR positive cell from Mex3a WT (left) and Mex3a cKO (right) immunofluorescence experiment described in (B). n = 1101 WT and 1278 Mex3a cKO OR positive cells from three biological replicates of four-week-old littermates. Statistics, Wilcoxon rank sum test. **D.** Representative Airyscan image of immunofluorescence for Arl13b in ciliary basal bodies. Scale bar, 2 µM. Magenta arrows highlight examples of basal bodies at origin of cilium projections. **E.** Quantification of Arl13b normalized integrated intensity within each dendritic knob using radial profile analysis. Radius of each knob normalized to one micron. n = 81 Mex3a WT dendritic knobs (blue) and 161 cKO dendritic knobs (yellow). Statistics, Fisher’s Z (1925), z = 19.313, p-value = 0.0000. **F.** Representative scanning electron microscopy image of OSN dendritic knob with cilium projections in Mex3a WT (left) and cKO (right). Scale bar, 1µM. **G.** Visual representation of basal body locations for each cilium projection over the dendritic knob for Mex3a WT (left) and cKO (right). All knob radii were normalized to 1 µM. **H.** Quantification of cilium projection distances from the center of the dendritic knob. n = 41 Mex3a WT dendritic knobs (blue) and 72 cKO dendritic knobs (yellow). Statistics, Wilcoxon rank sum test. **I.** Representative immunofluorescence image of MOE tissue Tgn46 in magenta, Mor28 OR in green, DAPI in blue. Scale bar 25 µM. **J.** Violin plot quantifying polarity of OSNs in Mex3a WT (left) and cKO (right). Polarity is quantified by measuring the angle between a line through the apex of Tgn46 and OR staining and the basal/apical axis of the epithelium where that cell resides (see diagram). n= 408 cells from WT, 328 from Mex3a cKO, 5 biological replicates each genotype at 4 weeks old. Statistics Wilcoxon rank sum test. **K.** Representative spinning disk confocal deconvolved SoRA live cell images of a HEK293T cell transiently transfected with mStayGold-UtrCH (green) and mCherry or Mex3a-mCherry (magenta). Scale bar, 10 µM. For all statistical tests in this figure: P<0.05 = *, P<0.01=**, P<0.001=***.

One explanation for reduced cilia-protein expression is that cilia are not properly formed in Mex3a cKO OSNs. To test whether cilia are present and properly formed in Mex3a cKO neurons, we performed immunofluorescence and high-resolution microscopy for the OSN cilia marker Arl13b^16^. In WT mice, Arl13b marks circles delineating each OSN dendritic knob and smaller circular structures marking basal bodies of each cilium projection (Figure 1D and S1F). Our imaging analysis reveals that in Mex3a cKO mice, the radial organization of basal body structures around the dendritic knob is not preserved (Figure 1D and E). Recent work unveiled dynamic organization of basal body positions as neurons mature^17^. To test whether the aberrant Mex3a cKO ciliary structures originated from immature OSNs, we used markers of immature neurons (Ngn1-GFP) and mature neurons (OMP-ires-GFP) to determine whether Arl13b signal colocalized with immature or mature neurons. In both Mex3a WT and cKO conditions, Arl13b colocalized with mature neuron dendritic knobs (Figure S1H), and not with immature neurons. This result confirms that defects observed in cilia structure originate from mature neurons in the Mex3a cKO context. Mex3a cKO mature, OMP-ires-GFP positive neurons harbor cilia with radial organization reminiscent of the pattern described^17^ in immature OSN cilia.

To further visualize OSN cilia structure in an antibody-independent way, and to confirm defects observed by immunofluorescence, we performed scanning electron microscopy (SEM) (Figure 1F). OSN cilia are present in both Mex3a WT and cKO mice, however, the cilium projections are arranged differently on the dendritic knobs in the two genotypes (Figures 1G-H). This spatial organization analysis demonstrates that WT cilia adopt a radially organized pattern around the dendritic knob, whereas cilia in Mex3a cKO mice display a largely random spatial distribution. Taken together, these microscopy data reveal that Mex3a is required for proper structure and patterning of dendrites in mature OSNs.

Given the links between cilia and planar cell polarity^18–20^, we wondered if polarity of olfactory sensory neurons was perturbed in Mex3a cKO mice. We performed immunofluorescence for Tgn46, a Golgi marker that is oriented on the apical side of the mOSN, along with four ORs (Mor28, P2, C7, and M71). We observed higher variability in the orientation of Mex3a cKO neurons within the tissue, including some cells oriented 180 degrees toward the basal side of the tissue instead of the apical side (Figure 1I-J). Our data demonstrate that in addition to cilia structure, cell polarity is disrupted in Mex3a cKO OSNs.

As cytoskeleton organization has known roles in cilia structure, planar cell polarity, and protein trafficking, we turned to an *in vitro* system to test whether increasing Mex3a levels affected cytoskeleton organization. We co-transfected HEK293T cells with either mCherry or Mex3a-mCherry and the filamentous actin marker mStayGold-UtrCH^21^. In HEK293T cells with mCherry, mStayGold-UtrCH readily marks filopodia/lamellipodia and reveals irregularly shaped HEK293T cells (Figure 1K and S1G). Upon co-transfection with Mex3a-mCherry, on the other hand, HEK293T cells are circular and lack the characteristic protrusions that denote lamellipodia. This rounded shape lends support to Mex3a’s role in cell shape, polarity, and cytoskeleton organization.

The data presented for loss of Mex3a in OSNs and increased Mex3a in HEK293T cells indicate that Mex3a regulates the interrelated processes of cytoskeleton organization, cilia structure, and membrane-protein synthesis or trafficking.

### Mex3a knockout leads to altered differentiation outcomes in the olfactory epithelium

Mex3a is expressed before and during activation of the Unfolded Protein Response (UPR) during OSN differentiation. The UPR signaling pathway is activated in several differentiation paradigms and facilitates translation and trafficking of membrane-bound proteins as cells become specialized^2,22^. In OSNs, UPR is required for maturation^23,24^. We previously showed that UPR transcription factor Atf5 and Atf5 downstream gene targets are more highly expressed in Mex3a cKO neurons^8^, and yet here we report deficits in mature neurons related to membrane-bound proteins, raising the possibility that UPR is not “resolved” in Mex3a cKO neurons. To test this, we turned to a previously published report which characterizes the transcriptional profiles of OSNs after UPR^25^. OSNs activate UPR to varying levels, depending on which OR is expressed, and mature neurons can be delineated into “high ER stress” and “low ER stress” neurons with distinct gene expression patterns^25^. Using single cell RNA-Seq (scRNA-Seq), we find that Mex3a cKO mature OSNs express both high and low stress ORs (Figure S2A). We used a module score analysis^26^ of 177 genes that were enriched in low stress WT OSNs and 941 high stress WT OSN enriched genes (Figure S2B). With this strategy we compared how Mex3a cKO low or high stress neurons expressed ER stress-specific genes compared to WT littermate control neurons. Curiously, ER stress genes were not upregulated to the same degree in Mex3a cKO mOSNs compared to WT littermate mOSNs (Figure S2C). Of note, known markers of mOSNs were expressed at the same or higher levels at the RNA level in Mex3a cKO mOSNs, revealing that reduced expression of ER stress genes was not due to Mex3a cKO mOSNs “arresting” at an earlier stage of differentiation (Figure S2D). One potential explanation for these results is that the ER stress response is not properly resolved in Mex3a cKO neurons.

We sought to clarify these seemingly contradictory findings: Mex3a cKO immature neurons express higher levels of Atf5 and Atf5 targets, but Mex3a cKO mature neurons do not properly upregulate signature genes that define high or low ER stress outcomes. We used proteomics and performed mass spectrometry on fluorescence activated cell sorted (FACs) populations of mature OSNs to identify 6026 protein groups across WT and Mex3a cKO samples (Figure 2A, Supplemental File 1). To gain insight into the types of proteins that are differentially abundant in the two genotypes, we performed Gene Ontology analysis on proteins that exhibited a log fold change (Mex3a cKO – WT) less than -0.58 (321 proteins, Figure 2B) or greater than 0.58 (165 proteins, Figure 2C). This analysis reveals that proteins related to ribosome biogenesis, protein trafficking, and lipid metabolism (required for cell membrane biogenesis) exhibit reduced protein levels in Mex3a cKO mature OSNs. On the other hand, proteins related to detoxification, processing amino acids, and cellular metabolism for energy production were measured at higher levels in Mex3a cKO mature OSNs than in control mature OSNs. We generated high confidence lists of significantly reduced and increased proteins in Mex3a cKO mature OSNs (Figures 2D and E). Significantly reduced proteins in Mex3a cKO cells were related to ciliogenesis or cilia function (Cfap20, Cngb1, Lrguk, Poc1b, Tubgcp3, and Rabl3), protein trafficking and vesicle dynamics (Ap5m1, Myo5b, Sec11a, and Use1), ribosome biogenesis and translation (Elp3, Nol9, and Rpl13a), and lipid metabolism, required for cell membrane biogenesis (Abca3, Agpat4, Hsd17b11, Nmt1, Ppm1l, and Srebf2). The functions of these dysregulated proteins support the phenotypes we observed in Mex3a cKO mature neurons (Figure 1).

**Figure 2:**
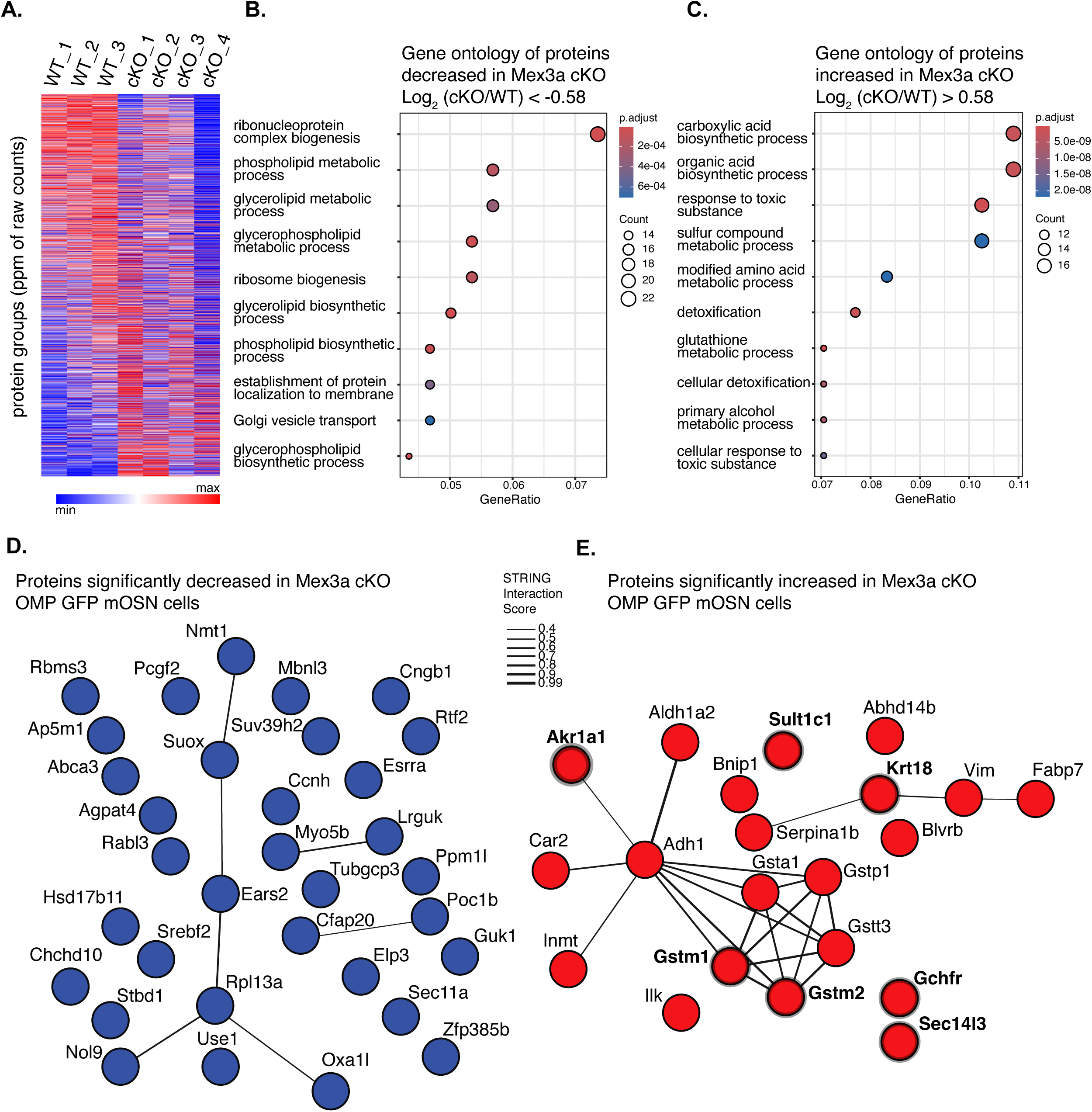
Mex3a knockout leads to altered differentiation outcomes in the olfactory epithelium. **A.** Heatmap generated with Morpheus Broad software of normalized protein group quantities for 6026 proteins (y-axis) measured by Mass Spectrometry across 7 samples (x-axis). Proteins were quantified from three replicates of OMPiresGFP+ sorted cells from Mex3a WT littermate controls, and four replicates of OMPiresGFP+ sorted cells from Mex3a cKO mice. **B.** Gene Ontology Analysis from 321 proteins found to be depleted from Mex3a cKO mature OSNs (LFC < -0.58). **C.** Gene Ontology Analysis from 165 proteins found at higher levels in Mex3a cKO mature OSNs (LFC > 0.58). **D.** STRING representation of proteins which exhibit a LFC (Mex3a cKO – WT) < -0.58, Student’s t-test p-value < 0.05, and raw protein counts in all 7 samples to ensure sufficient coverage. **E.** STRING representation of proteins that are increased in Mex3a cKO samples (LFC > 0.58), Student’s t-test p-value < 0.05, and raw protein counts in all 7 samples. Proteins with thick borders are markers of sustentacular cells as determined by scRNA-Seq from MOE (data from^8^).

Multiple proteins that were significantly increased in Mex3a cKO neurons were related to detoxification (Akr1a1, Gsta1, Gstm1, Gstm2, Gstt3, Sult1c1). Intriguingly, detoxification and processing of xenobiotic compounds are primary roles of sustentacular cells^27^, which are progeny of the globose basal cells where Mex3a is highly expressed^8^. Using our scRNA-Seq dataset, we identify markers of sustentacular cells and found that seven sustentacular markers were among significantly increased proteins in Mex3a cKO mature neurons (Figure 2E). Intriguingly, when comparing transcriptional differences between Mex3a cKO and WT sustentacular cells by scRNA-Seq, we observe that markers of mature OSNs are upregulated in Mex3a cKO sustentacular cells (Figure S2E). In the absence of Mex3a, mature neurons and sustentacular cells are still present and are morphologically similar to WT cells, but they exhibit “lineage mixing” at the transcriptional/protein levels (Figure S2F). These findings uncover a striking role for Mex3a in enforcing lineage fidelity, selectively repressing sustentacular programs in neurons and neuronal programs in sustentacular cells.

Our molecular analysis of Mex3a cKO neurons reveals that while mature markers are expressed, cells exhibit deficits in maturation including reduced expression of genes that are downstream of UPR. This combined with reduced levels of proteins involved in trafficking, translation, and membrane dynamics all point towards a possible defect in “resolving” UPR to allow for proper membrane-bound protein expression. In addition, our proteomics and scRNA-Seq data reveal that in the absence of Mex3a, the neuronal and sustentacular lineages are less well defined and instead genes from the other lineage become ectopically expressed. Taken together our data support a role for Mex3a in establishing crucial expression paradigms in immature cells that support proper neuronal (and sustentacular) differentiation.

### Putative RNA targets of Mex3a uncovered by *in vivo* HyperTRIBE

Mex3a harbors two KH-RNA binding domains as well as intrinsically disordered domains (IDRs) (Figure 3A), which, among other functions, facilitate membrane-less organelles to form in cells. Proteins rich in IDRs are known to be enriched in cytoplasmic granules involved in RNA degradation, modification, storage, and translation^28^. To visualize Mex3a subcellular localization in live cells, we co-transfected HEK293T cells with Mex3a-mCherry and ER marker GFP-Sec61b^29^ constructs (Figure 3B). We observe Mex3a-positive puncta, sometimes in close association with the ER, reminiscent of cytoplasmic granules. These imaging data lend support to Mex3a’s role as an RNA binding protein, residing in cytoplasmic granules that likely regulate RNA. We sought to identify Mex3a’s RNA targets that could be subject to post-transcriptional regulation in immature OSNs.

**Figure 3:**
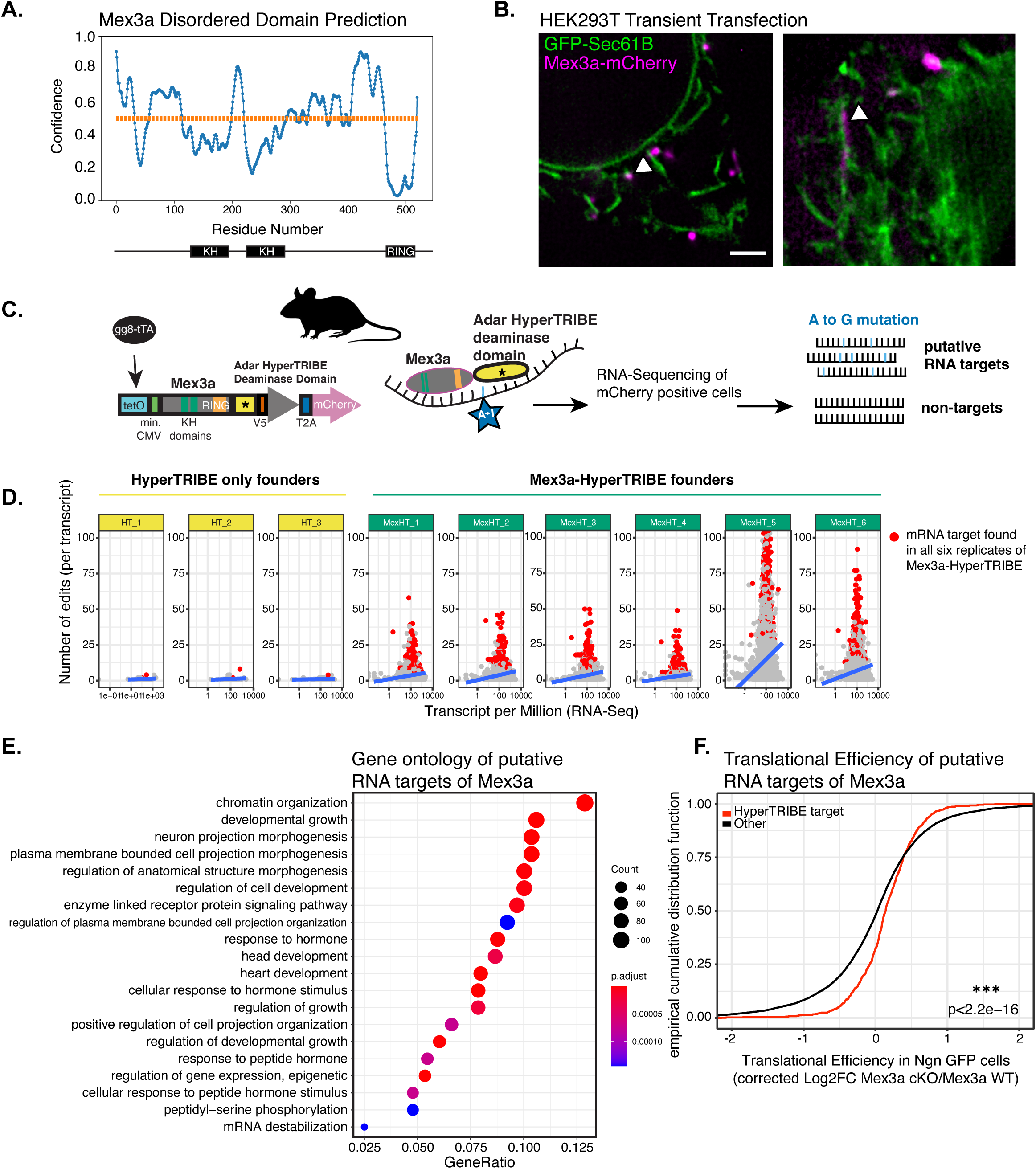
Putative RNA targets of Mex3a uncovered by *in vivo* HyperTRIBE. **A.** Disordered domain prediction for Mex3a protein, as determined using PrDOS server^115^ **B.** Live cell imaging of HEK293T cells transiently transfected with GFP-Sec61b and Mex3a-mCherry. Images acquired with high resolution SoRA microscope and deconvolved using FIJI plugin Microvolution. Scale bar, 2 µM. **C.** Design of transgenic allele used to find RNA targets of Mex3a *in vivo* in the olfactory epithelium. The mouse Adar deaminase domain with a HyperTRIBE point mutation (E957Q) was cloned downstream of the Mex3a gene, followed by a V5 tag, t2a and mCherry. The construct is induced by a tetO with minimal CMV promoter, driven by gg8-tTA or OMP-tTA in the MOE. After sorting mCherry positive and negative cells from the MOE, RNA-Seq is conducted and A→I mutations (read out as G after sequencing) are used to infer RNA targets. **D.** Scatterplot comparing the number of editing sites per transcript on the y-axis (set to 100 for each for comparison) to expression level (TPM) of that transcript on the x-axis. mRNAs shown in red were found in all 6 mCherry+ RNA-Seq libraries from six different tetOMex3a-HyperTRIBE transgenic founders. While 1-2 of these mRNAs were observed in each HyperTRIBE-only sample, they were edited to a lesser extent and were not the same mRNAs across all 3 of the HyperTRIBE-only samples. **E.** Gene Ontology analysis of putative HyperTRIBE RNA targets of Mex3a after driving the construct in mice with the gg8-tTA driver. Targets were given a weighted score based on number of editing sites and how frequently the same RNA was edited across replicates. The RNAs with scores from 0.85 to 1 were considered putative targets. **F.** ECDF graph comparing the translational efficiency scores in Ngn1-GFP sorted cells^8^ of HyperTRIBE targets compared to non-HyperTRIBE RNAs. Statistics: Two-sample Kolmogorov-Smirnov test. For all statistical tests in this figure: P<0.05 = *, P<0.01=**, P<0.001=***.

Proteomes of Mex3a cKO neurons reveal global outcomes of loss of Mex3a, but do not provide direct targets of Mex3a that would explain Mex3a’s mechanism of action. To identify putative RNA targets *in vivo*, we adapted the HyperTRIBE technique described in drosophila^30^ to mouse (Figure 3C and S3A). We first tested and concluded that mouse Adar deaminase domain was the most active, compared to paralogs Adarb1 and Adarb2 (see Materials and Methods). Next, we found that the E448Q HyperTRIBE mutation identified in drosophila is conserved in mouse (E957Q). We made transgenic mice that harbored a tetO-HyperTRIBE construct (three founders) or a tetO-Mex3a-HyperTRIBE construct (six founders). We first bred these transgenic mice to gg8-tTA^31^ and OMP-ires-tTA^32^ mice to express the transgene in both immature and mature OSNs. After RNA-Seq of FAC sorted transgene expressing cells, we observed high levels of Adar expression in progeny from all founders, and high levels of Mex3a in tetO-Mex3a-HyperTRIBE progeny (Figure S3B). RNA editing was quantified across all samples (Figure S3C). High editing rates in tetO-Mex3a-HyperTRIBE samples compared to tetO-HyperTRIBE samples reveal that deamination by the Mex3a-HyperTRIBE allele is specific and not the result of off-target deamination by the HyperTRIBE domain alone. Furthermore, we found high concordance for which RNAs exhibited editing across all six tetO-Mex3a transgenic founders (Figure 3D). To quantify editing strength, we derived a HyperTRIBE score: a weighted average that combines the number of edits per RNA with how consistently those RNAs were edited across biological replicates originating from different transgenic founders. RNAs edited frequently and reproducibly received higher scores. To our knowledge, our strategy is the first use of genetically encoded inducible HyperTRIBE constructs in mice.

After determining that our constructs led to specific and reproducible RNA targets *in vivo*, we repeated the experiment in gg8-tTA; tetO-Mex3a-HyperTRIBE mice. Mex3a is endogenously expressed in gg8-tTA positive cells, and direct targets of Mex3a should be present in these immature OSNs. We generated HyperTRIBE scores for all RNAs identified with editing in our gg8-tTA; Mex3a-HyperTRIBE sorted cell RNA-Seq and identified putative RNA targets of Mex3a (See Materials and Methods, 932 RNAs, Supplemental File 2). Gene Ontology analysis of these putative targets reveals transcripts involved in chromatin organization, neuron projection organization, and response to hormone (Figure 3E). Of note, two top putative RNA targets are related to Wnt signaling (Wnt receptor Fzd3 and Usp34, a ubiquitin hydrolase activator of the Wnt pathway). These and other Wnt-related RNA targets of Mex3a can explain planar cell polarity defects observed in Mex3a cKO mature OSNs.

Given Mex3a’s role as a repressor in C. elegans^3,33^, we wondered if loss of Mex3a would cause translational de-repression of these putative Mex3a RNA targets in MOE. To test this, we examined our previously published translational efficiency scores from Ngn1-GFP (immature OSN) sorted Mex3a cKO and WT mice^8^ to compare putative HyperTRIBE targets to non-targets. We observe a significant increase in translational efficiency of putative RNA targets in Mex3a cKO Ngn1 cells, suggesting that Mex3a has a repressive effect on the translation of its RNA targets in the WT context (Figure 3F).

Our strategy uncovers Mex3a RNA targets *in vivo* in the MOE. We find putative targets with direct links to the phenotypes we observe in Mex3a cKO neurons, including Wnt signaling genes and genes related to neuron projection organization. Analysis of our previously published ribosome profiling data in sorted immature neurons shows that RNA targets identified by HyperTRIBE are translationally derepressed in Mex3a cKO immature neurons, providing evidence that Mex3a is a translational repressor.

### Mex3a Ubiquitin targets identified in MOE are implicated in trafficking and translation

In addition to RNA binding domains, Mex3a harbors a RING E3 ubiquitin ligase domain which we hypothesized would be essential for its function. To understand what proteins are ubiquitin targets of Mex3a, we adapted UbiFast^34^ to our mouse Mex3a cKO model. We reasoned that Mex3a ubiquitin targets would be less ubiquitinated in Mex3a cKO samples compared to littermate controls. We isolated and lysed whole MOE, performed trypsin digest and immunopurification of the K-ε-GG epitope that is generated upon trypsin digest. This provides peptide resolution and enrichment of ubiquitin containing proteins, and provides a dataset illuminating which proteins are post-translationally ubiquitinated in the MOE (Supplemental File 3). We next compared the ubiquitinomes of Mex3a cKO MOE to Mex3a WT MOE and identified 61 putative Mex3a ubiquitin targets (Figure 4A, Supplemental File 4), including proteins found in mitochondria, cytoskeleton, and endomembrane cellular components. We note that not all changes in ubiquitination necessarily indicate direct Mex3a substrates. Some proteins may exhibit altered ubiquitination as an indirect consequence of the broader perturbations caused by Mex3a cKO in the MOE. We were intrigued to find putative Mex3a targets with known roles in UPR (Gap43, marker of iOSNs by scRNA-Seq, ubiquitinated at K114, Cebpg^23^ at K113, and Rad23b at K36), translation (Serbp1, ubiquinated at K68, and Rps7 at K10), and protein trafficking (Abcf3 ubiquitinated at K296 and/or K302, Epn1 at K128, Pip5k1a at K88, Rnf13 at K233, Sec14l1 at K71, Sec14l3 at K117, Sec31a at K689 and/or K728, and Stx12 at K82).

**Figure 4:**
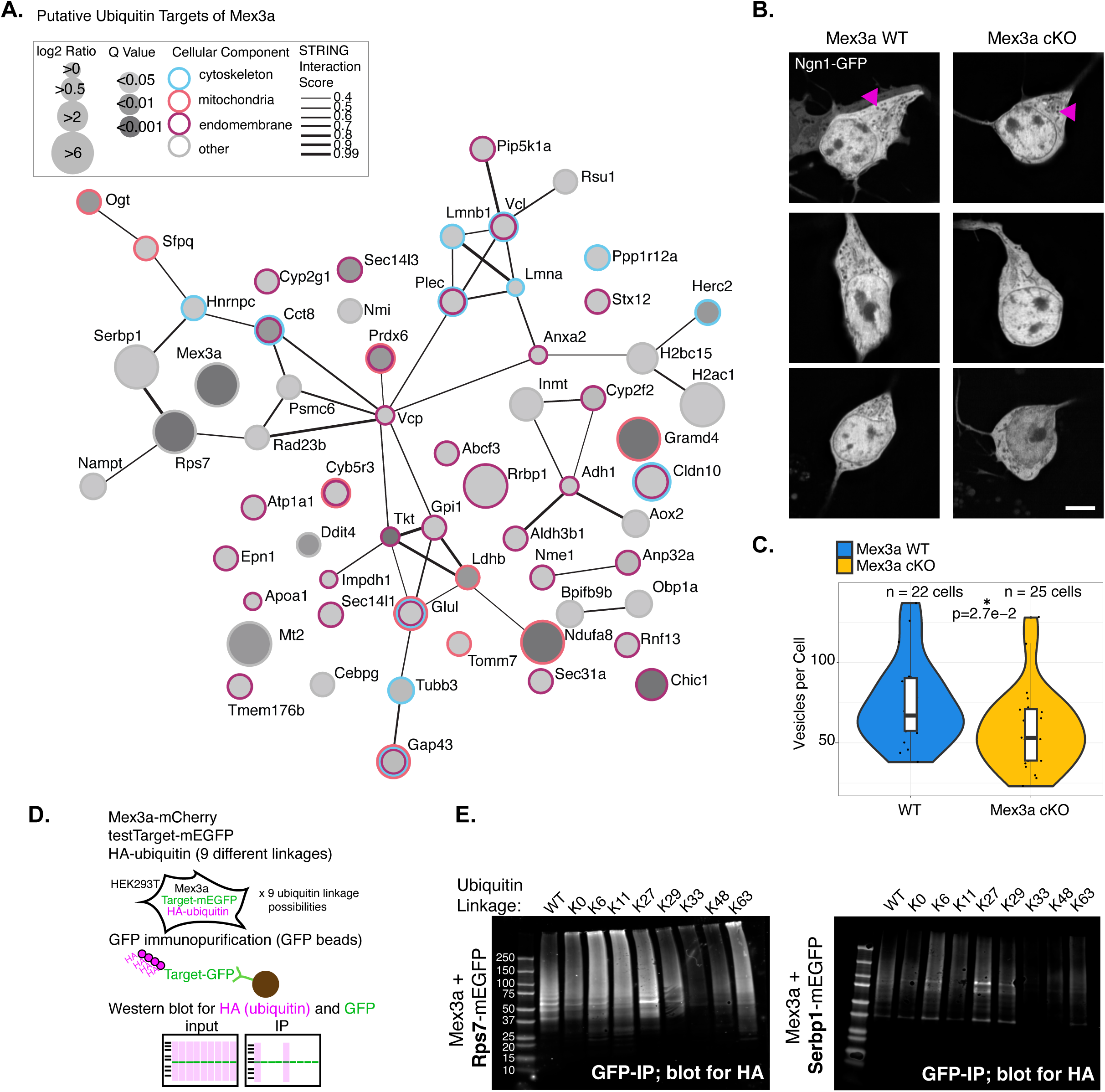
Mex3a Ubiquitin targets identified in MOE are implicated in trafficking and translation. **A.** STRING representation of putative ubiquitin targets of Mex3a. Ubiquitinated peptides were enriched from Mex3a WT and cKO MOE and proteins with significantly higher ubiquitination in the WT context were considered putative targets of Mex3a. Size of circle relates to log2 ratio of enrichment in WT compared to Mex3a cKO, and opacity of circle reflects Q value. Proteins are circled with blue (cytoskeleton), pink (mitochondria), or magenta (endomembrane) rings to represent cellular localization. STRING interactions depicted as weighted lines with more confident interactions shown as thicker lines. **B.** Representative live cell images from cultured OSNs, Mex3a WT and cKO; Ngn1-GFP. Magenta arrows point towards examples of vesicles measured in analysis. **C.** Quantification of number of vesicles per cell, n = 22 WT cells, 25 Mex3a cKO cells, two biological replicates each genotype. Statistics: Wilcoxon rank sum test. **D.** Experimental design to test which ubiquitin linkage is conferred onto putative ubiquitin targets Serbp1 and Rps7. Each HA tagged ubiquitin construct harbors lysine→arginine mutations to allow only one ubiquitin-linkage, except for the WT-ubiquitin construct which has all lysines intact. HEK293T cells are transiently transfected, and mEGFP construct is immunopurified with GFP-magnetic beads. Input and immunopurified samples are run by SDS page for each condition, and Western blot is performed for HA and GFP (Figures 4E and S4A-C). **E.** Western blot for HA after immunopurifying mEGFP tagged constructs (Rps7, left, Serbp1, right). Molecular weight of ladder markers shown on left Western. See Figure S4A-C for additional controls. For all statistical tests in this figure: P<0.05 = *, P<0.01=**, P<0.001=***.

As multiple putative ubiquitin targets of Mex3a were involved in trafficking, and given the phenotypes and proteomics data revealing deficits in membrane-bound protein transport, we quantified cytoplasmic vesicles, essential for membrane-protein trafficking, in living immature neurons. To quantify vesicles, we bred the Ngn1-GFP allele into our Mex3a cKO mice to mark immature neurons and to take advantage of the fact that GFP does not integrate into cytoplasmic, membrane-coated organelles and vesicles. We cultured neurons and imaged living cells in the GFP channel. After quantification of small, circular structures with no GFP (see Materials and Methods), we count fewer cytoplasmic vesicles in Mex3a cKO neurons compared to Mex3a WT littermate controls processed in parallel (Figure 4B and C). Taken together with our proteomics data and phenotypic characterization, these data further support deficits in protein trafficking.

Our UbiFast experiment gives peptide resolution of where on a protein a ubiquitin moiety can be found. However, as trypsin digests the ubiquitin protein just two amino acids after the modified lysine, ubiquitin branching information is lost. Identifying which ubiquitin linkage Mex3a adds clarifies its function, as specific polyubiquitin chains can mark substrates for proteasomal degradation, alter signaling, or regulate other non-proteolytic processes. To test which ubiquitin chain Mex3a confers on substrates, we co-transfected three constructs into HEK293T cells: 1) Mex3a-mCherry 2) one putative ubiquitin target (either Serbp1-mEGFP or Rps7 mEGFP) and 3) one of nine different HA-tagged ubiquitin constructs (Figure 4D). Except for the WT ubiquitin positive control, each of the eight additional ubiquitin constructs had lysine-to-arginine mutations that render only one type of ubiquitin linkage possible. Enrichment of that ubiquitin construct after IP suggests that the target protein is modified with that specific type of ubiquitin chain. Input Westerns demonstrated that all constructs were well expressed in HEK293T cells except for HA-K33-ubiquitin (Figure S4A-C), thus we cannot conclude whether Mex3a confers K33-ubiquitination or not. We immunopurified mEGFP-tagged constructs from protein lysates, then used Western to blot for HA-tagged ubiquitin constructs (Figure 4E). Our results reveal strongest enrichment for K27 ubiquitin, reported to modulate protein activity, on both Serbp1 and Rps7 when co-transfected with Mex3a. This experiment provides evidence that our UbiFast experiment is capable of uncovering primary ubiquitin targets of Mex3a. In addition, our analysis in HEK293T cells suggests that Mex3a K27-ubiquitinates Serbp1 and Rps7, supporting a non-proteolytic role for Mex3a ubiquitination. K27 ubiquitination could instead lead to increased activity of targets^35^, regulate phosphorylation or dephosphorylation of targets^36^, or affect substrate binding to targets^37^.

### Mex3a levels are associated with changes in Serbp1 and phosphorylated eEF2 levels in ribosomes

Given our previous finding that Mex3a associates with ribosomes^8^ in addition to our current results suggesting translational de-repression of RNA targets in Mex3a cKO neurons, coupled with putative ubiquitin targets involved in translation, we wondered what role, if any, Mex3a played in translation. To explore this, we tested how both loss of Mex3a and increase in Mex3a levels affected monosome and polysome area and recruitment of translation factors. We also used polysome fractions to explore changes in mRNA abundance in ribosome fractions from Mex3a cKO MOE tissue.

Sucrose gradients and ultracentrifugation of MOE tissue lysis from WT and Mex3a cKO mice were used to separate ribosomes by density and perform polysome profiling (Figure 5A). Our polysome analysis shows that Mex3a cKO MOE has a higher polysome to monosome ratio and lower monosome area than WT littermate MOE (Figures 5B and S5A). This could imply that Mex3a has a role in repressing translation initiation and/or enhancing translational elongation.

**Figure 5:**
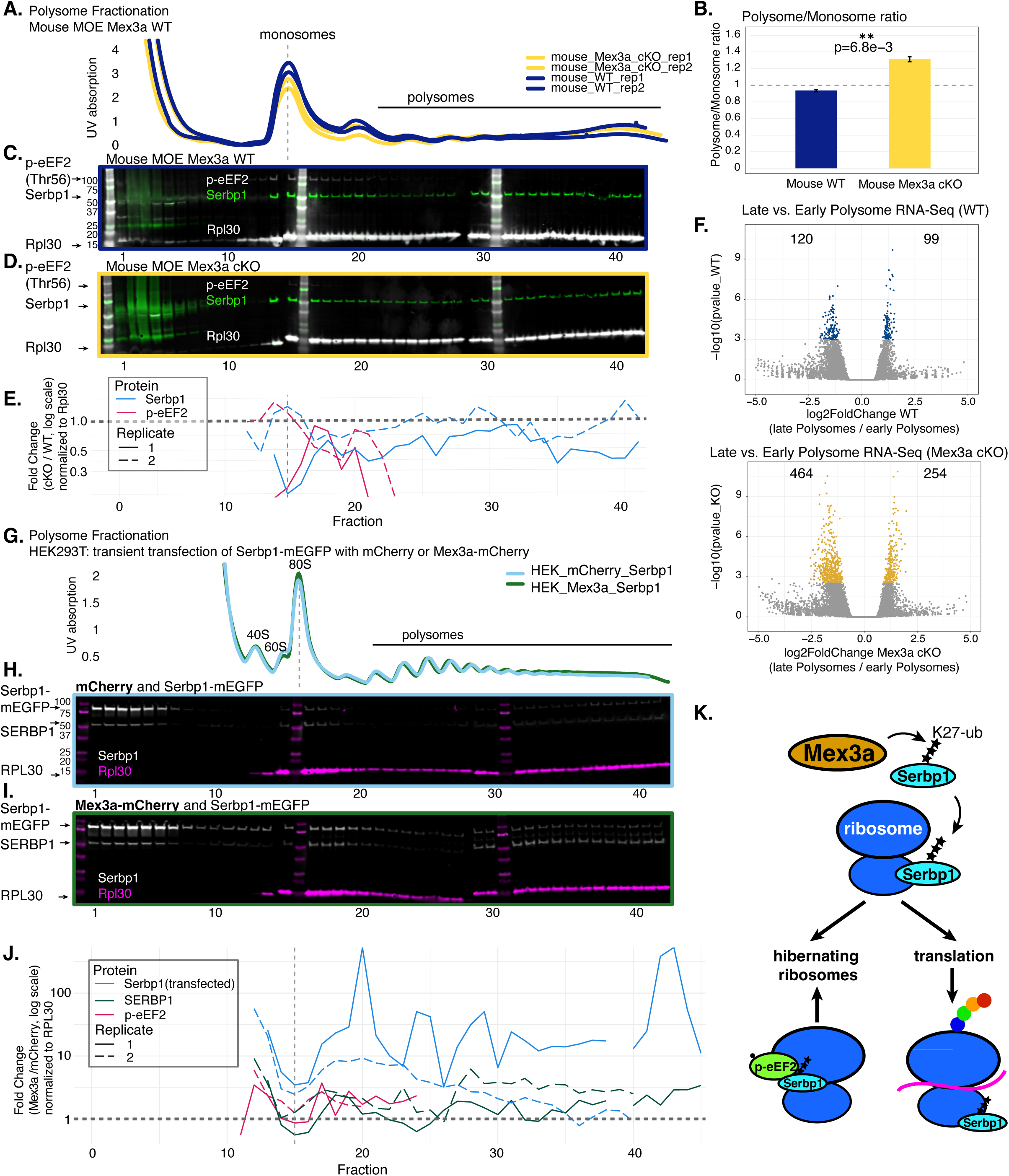
Mex3a levels are associated with changes in Serbp1 and phosphorylated eEF2 levels in ribosomes. **A.** Polysome profiling from Mex3a WT or cKO MOE tissue, two biological replicates for each genotype consisting of 10 pooled MOE for each replicate, 20 MOE for each genotype. Figure generated with QuAPPro^116^. **B.** Polysome to Monosome ratios for WT and Mex3a cKO samples. Area of monosome and polysomes measured using QuAPPro, and ratio quantified by dividing the area of polysomes by area of monosomes. Error bars, SEM. Statistics, Student’s t-test. Data from two biological replicates for each genotype. **C.** Representative Western blot for endogenous mouse eEF2 (phosphorylated at Thr56), Serbp1, and Rpl30 in fractions from Mex3a WT. Molecular weight of ladder markers shown applies to (C) and (D). **D.** same as (C) but for Mex3a cKO polysome fractions. **E.** Quantification of Western blot for Serbp1 and p-eEF2 presented as fold change of Mex3a cKO over WT, each fraction normalized to Rpl30 value from the same fraction. Data are plotted by fraction on the x-axis, with a thin dotted line where the monosome peak is observed by polysome profiling. Thick dotted line at y=1 to show positive (above the line) or negative (below the line) fold change compared to Mex3a WT. Data presented from two replicates each genotype. **F.** Volcano plots showing differentially abundant RNAs in late (fractions 35-42) versus early (factions 27-34) for Mex3a WT (top) and Mex3a cKO (bottom) identified using DESeq2, results calculated with lfcThreshold = 0.5849 for more stringent statistical test. log10 p-value is plotted on y-axis, but transcripts were considered significant only if adjusted p-value was < 0.05. **G.** Polysome profile of HEK293T cells transiently transfected with Serbp1-mEGFP and Mex3a-mCherry (dark green line) or Serbp1-mEGFP and mCherry (light blue line). Profiles were aligned at peak of monosome (dotted line). Figure generated with QuAPPro. Two replicates for each condition. Five 10cm dishes pooled per condition and replicate. **H.** Forty-two fractions were collected and RPL30 and Serbp1-mEGFP/SERBP1 levels were determined by Western blot for each fraction from the mCherry and Serbp1-mEGFP transfection. Note that bands at the expected sizes for human SERBP1 and mouse Serbp1-mEGFP are seen after blotting with the Serbp1 antibody. Molecular weight of ladder markers shown applies to (H) and (I). **I.** Representative Western blot for Serbp1 and RPL30 for polysome fractions from Mex3a-mCherry and Serbp1-mEGFP transfection. **J.** Fold change (Mex3a-mCherry transfection/mCherry transfection) of human SERBP1, mouse Serbp1 and p-eEF2 Western signals, normalized to RPL30. Thick dotted line at y=1 to show positive (above the line) or negative (below the line) fold change relative to mCherry control. Data presented from two replicates each of five pooled 10cm cell culture dishes per condition. **K.** Model depicting Mex3a conferring K27 ubiquitin on Serbp1 which promotes its recruitment to ribosomes and modulates hibernating ribosomes and/or translational initiation/elongation. For all statistical tests in this figure: P<0.05 = *, P<0.01=**, P<0.001=***.

Mex3a ubiquitin target Serbp1 (Figures 4A and E) is known to reside in cytoplasmic granules and associate with ribosomes^38,39^. In addition, upon phosphorylation of eEF2 at threonine 56, Serbp1 has been reported to translocate into the A and P sites of the ribosomal mRNA channel. This has the dual role of blocking active translation and preserving ribosomes in a hibernating state during times of cellular stress^40–42^. We tested whether modulating levels of Mex3a would affect Serbp1 and p-eEF2 recruitment to ribosomes by performing Western blot on polysome profiling fractions.

Our polysome and Western analysis reveals reduced Serbp1 at both monosomes and polysomes, as well as reduced p-eEF2 at monosomes in Mex3a cKO MOE (Figure 5C-E). Importantly, as no antibody is available for ubiquitinated or phosphorylated Serbp1, quantification of total levels of Serbp1 may not reflect post-translationally modified Serbp1 levels, which could have important functional consequences on translation^43^.

To test whether we could identify functional changes in translation rates upon loss of Mex3a, we performed RNA-Seq on pools of early (fractions 27-34) and late (fractions 35-42) polysomes. When comparing RNA-Seq of polysome fractions, we observed that the Mex3a cKO samples exhibited a larger number of genes with significant changes between late polysome fractions and early polysome fractions than the WT samples (Figure 5F, Supplemental File 5). These results suggest greater variability in mRNA abundance across polysome fractions in the Mex3a cKO context, possibly due to changes in translation rates or efficiency. Formal tests comparing genotype context across fractions did not detect a significant interaction, possibly due to small sample size, or as the analysis was performed using whole MOE tissue, the effects on translation in Mex3a-expressing cells is diluted by translation of those mRNAs in other cell types. Further investigation is required to determine whether Mex3a regulates translation rates or efficiency in Mex3a-expressing cells or their progeny.

We wondered how increased Mex3a levels would affect monosome area and recruitment of Serbp1 and p-eEF2 to ribosomes. To test this, we turned to our heterologous HEK293T transient transfection system, transfecting mouse Serbp1-mEGFP with either mouse Mex3a-mCherry or mCherry contructs. After polysome profiling followed by Western blot, we find both endogenous human MEX3A and transfected mouse Mex3a-mCherry in monosome and polysome fractions, with particular enrichment in fractions corresponding to the 40S small subunits (Figure S5B). This result is supported by our previous IP-Mass Spectrometry results in mouse showing that Mex3a interacts with ribosomes^8^. Our polysome profiles revealed an increase in monosome area in HEK293T cells transfected with Mex3a and Serbp1 as compared to mCherry and Serbp1 transfected cells (Figures 5G, S5A and S5C). Both endogenous human SERBP1 and transfected mouse Serbp1-mEGFP were detected by the Serbp1 antibody, and we found that when Mex3a-mCherry was co-transfected, Serbp1 and SERBP1 levels increased in ribosomes compared to the mCherry transfection condition (Figure 5H-J). Endogenous human p-eEF2 levels also increased around monosome fractions when Mex3a levels were higher (Figures 5J and S5D).

Taken together our loss and gain of Mex3a in mouse and HEK293T cells, respectively, reveals that Serbp1 and p-eEF2 incorporation into ribosomes is positively correlated with Mex3a levels. In the absence of Mex3a, Serbp1 and p-eEF2 are not recruited to ribosomes as efficiently, likely leading to changes in polysome to monosome ratios and less well-regulated translation *in vivo.* Our ubiquitin and polysome profiling data lead to a model (Figure 5K) where Serbp1 is K27-ubiquitinated by Mex3a which leads to its recruitment to ribosomes. We predict that this affects p-eEF2 recruitment to ribosomes which would modulate hibernating ribosome levels and/or translation initiation/elongation. We propose that Mex3a ubiquitinates targets in immature OSNs where it is expressed, and these post-translational modifications have lasting effects on mature OSNs even after Mex3a is silenced. In addition, our data provide evidence that Mex3a translationally represses RNA targets, likely regulating the timing of their expression. Loss of Mex3a in the OSN lineage results in lineage inappropriate gene expression, as well as reduced translation/trafficking of membrane-bound proteins and neuron projection defects.

## Discussion

### Developmental defects in Mex3a cKO neurons

While in the MOE, Mex3a is exquisitely restricted to immature OSNs, this work uncovers previously unappreciated roles for Mex3a that ensure proper neuronal maturation. Curiously, the cells that exhibit the most striking phenotypes upon loss of Mex3a – reduced mRNA translation and trafficking of cell surface proteins, abnormal cilia structure, changes in planar cell polarity, and axon mistargeting^8^ are the fully differentiated, mature OSNs that would have ceased to express endogenous Mex3a. This lends support to the essential role that post-transcriptional and post-translational regulation plays in neuronal development^44^ and reveals that there is a “memory” of crucial regulation in immature cells that persists and has important consequences in mature cells.

At the molecular level, mature neurons from Mex3a cKO mice express markers of mature neurons, however, they exhibit increased expression of sustentacular cell markers relative to controls. Likewise, sustentacular cells from Mex3a cKO mice show increased levels of neuronal markers. These fascinating results attribute Mex3a to ensuring proper lineage delineation, silencing lineage-inappropriate genes as cells differentiate from globose basal cells to neurons or sustentacular cells. A recent report in moss reveals that lineage gatekeeping by post-translational regulators is a highly conserved method of ensuring proper differentiation^45^.

While in the MOE, loss of Mex3a does not change cell cycle dynamics^8^, Mex3a is perhaps most famous for its role in cell cycle^46–51^. Molecular targets of Mex3a that are related to cell cycle in other cell types might also be targets in OSNs, but instead of dictating cell cycle in OSNs, regulate related processes. Cytoskeleton organization, for example, especially regulation of centrioles required for ciliogenesis, is closely linked to cell cycle^52^. A recent study revealed that centrioles, which determine cilium position, are arranged over the dendritic knob in immature OSNs and migrate radially in mature OSNs^17^. Our data show that in Mex3a cKO mOSN dendritic knobs, this radial migration has not occurred. Could dysregulation of tubulin or other microtubule-related proteins account for this basal body migration defect? In support of this hypothesis, we find Tubb3 as a putative ubiquitin target of Mex3a, and microtubule nucleating protein Tubgcp3 is significantly depleted from Mex3a cKO mature OSNs at the protein level. In addition, previous work has shown that microtubule translation is selectively upregulated by increasing p-eEF2 at monosomes^53^. This supports our findings in the Mex3a cKO MOE which exhibits reduced p-eEF2 at monosomes coupled with phenotypes consistent with reduced microtubule translation and/or activity.

### Mex3a at the intersection of UPR, stress granules, and translational control during OSN differentiation

Our data indicate that Mex3a acts within UPR signaling. In Mex3a cKO immature OSNs, translation of UPR-activated transcription factor Atf5 is increased, and downstream targets of Atf5 are upregulated^8^. However, when considering the transcriptomes of mature OSNs, the outcome of UPR (high and low ER stress gene profiles^25^) are not properly upregulated in Mex3a cKO mature OSNs. This would suggest that UPR is not properly “resolved” if Mex3a is not present. Taken together these data provide evidence that Mex3a serves to repress UPR in immature OSNs where it is expressed, but serves an important role in “setting up” the cell for proper resolution of UPR in order to differentiate.

How might these seemingly incongruent phenotypes be explained? One possible explanation for reduced membrane-bound protein expression in mature OSNs, and apparent reduced ability to recover from UPR in absence of Mex3a, is a failure to properly traffic cell surface proteins. These proteins require cytoplasmic vesicles for transport from the ER to the Golgi and then to the plasma membrane. Our total proteomics and live cell imaging data in neurons reveal that vesicles and some vesicle trafficking proteins are significantly depleted from Mex3a cKO neurons. Previous work identified transcription factor Ddit3 as critical for establishing high and low ER stress gene profiles^25^. Could maintenance of these gene profiles be dependent on some feedback signal originating from cell surface proteins? If so, impaired trafficking of membrane-bound proteins could explain reduced expression of ER stress profile genes observed in Mex3a cKO mature OSNs.

A different model to test would be whether under conditions of cellular stress such as UPR, Mex3a is incorporated into ribosomes where it promotes poised translational complexes. Our polysome profiling experiments provide evidence that Mex3a promotes Serbp1 and p-eEF2 incorporation into ribosomes, which could lead to an increase in stalled, or “hibernating” ribosomes^38^. This might temporarily lead to reduced (or even stalled) translation, as we expect during ER stress, while protecting a fraction of ribosomes from ribophagy so that once UPR is resolved, translation could resume at normal levels. In the absence of Mex3a, reduced Serbp1 and p-eEF2 in monosomes might mean more ribosomes would be subject to ribophagy during ER stress, possibly leading to reduced ribosomes in mature OSNs. This model could explain the increase in translation of OR proteins that we report in immature OSNs^8^, as well as the reduced translation of OR proteins in mature OSNs presented here. Whether hibernating ribosomes would somehow be marked to translate cell-surface proteins, or whether they would be more general, all-purpose translational complexes would require further exploration.

Our data support previous work^54^ that Mex3a is a stress granule (SG) protein. Stress granules are membrane-less cytoplasmic granules that form dynamically in response to diverse cellular stresses and which have known roles in translational repression^55,56,57^ and neuronal development^58^. Stress granules can form as a result of unfolded proteins in the ER and PERK-mediated phosphorylation of eIF2α, which occurs in OSNs^24^. In addition, the ER is important in biogenesis of SGs^59^ and ER targeted mRNAs can be selectively recruited to SGs^60^. Our live cell microscopy in HEK293T cells reveals Mex3a-mCherry positive structures reminiscent of SGs, often in close association with the ER, as described previously^59^. Seminal work on Mex3a found it associated with Ago2^61^, which is incorporated in both P bodies and SGs. In addition to Ago2, we find other SG proteins to interact with Mex3a^8^ including Pabpc1, Atxn-2, Fmr1, Serbp1, Fam120a as well as Ddx, and Hsp family proteins^62–64^. We also report that HyperTRIBE putative RNA targets are translationally de-repressed in Mex3a cKO immature OSNs. This suggests that Mex3a has a repressive effect on translation of its RNA targets, which is consistent with SGs as sites for translationally stalled or repressed mRNAs. If Mex3a is important for SG formation or function during UPR, then loss of Mex3a could result in the defects related to translation that we observe in Mex3a cKO mature neurons.

As Mex3a is expressed before and during UPR signaling in immature OSNs, we hypothesize that UPR changes Mex3a activity, possibly to promote stalled translational initiation complexes at a critical time for translational control in the cell. We find evidence that Mex3a can be ubiquitinated at 9 different lysines in MOE: K27, K114, K120, K127, K209, K242, K294, K329, K454, and previous reports show that it is phosphorylated^65,66^. Post-translational modifications likely serve as a molecular switch, rapidly changing Mex3a’s activity, localization, or protein stability in response to changes in the cellular environment.

### Mex3a in health and disease

Mex3a is expressed in immature neurons and promotes neurogenesis^5,11,67,68^. In the highly regenerative intestinal crypt, Mex3a maintains a key reserve stem cell population through its regulation of Cdx2 and Lgr5^6,13,69,70^. Mex3a is also expressed in vascular tissues, promotes angiogenesis^71,72^, and plays a protective role against atherosclerosis^73^. Given its role as a stemness factor^74^, it is perhaps not surprising that Mex3a is increasingly recognized as a hallmark of metastatic cancers and cancers with poor outcomes. A growing number of studies has identified Mex3a as a prognostic biomarker of breast, ovarian, cervical, lung, soft tissue, hepatocellular, thyroid, colon, gastric, Wilms tumor, bladder, and pancreatic cancers^47,48,54,75–88^. This daunting list of sarcomas, carcinomas, and other cancers highlights the urgent need to understand the Mex3a regulome and molecular mechanism of action.

On a tissue level, Mex3a is reported to promote endothelial-to-mesenchymal transition (EMT), migration, and invasion, characteristic of metastasis^89–92^. On a cellular level, Mex3a has been linked to autophagy through interaction with and regulation by Ago2 and mir-126-5p^93–95^. On a molecular level, it is reported to be active in diverse signaling pathways including the Wnt/ß-catenin pathway^13,69,74,89,91,96–99^, the PIK3/AKT pathway^84,90,100–103^, NF-KB/Rig-I^104–107^, Hippo^108^ and RhoA/ROCK1^109^ pathways.

This study implicates Mex3a in cytoskeletal regulation which is one of the major targets of chemotherapies to treat cancer. We find that exogenous Mex3a expression in HEK293T cells forms more circular cells and blocks the formation of actin-rich lamellipodia/filopodia. Loss of Mex3a in olfactory sensory neurons leads to structural changes in dendrites, reduced trafficking of cell-surface proteins, and planar cell polarity defects, all of which are at least partially regulated by cytoskeletal proteins, especially microtubules^110–112^. Cytoskeletal dynamics regulate EMT and metastasis^113^, both of which have been previously linked with increased Mex3a expression in cancer cell types^89–92^.

Here we find that Mex3a can be found in association with ribosomes and appears to modulate translation factor localization to monosomes and polysomes. According to our model, an over-abundance of Mex3a, which is the case for many cancers, could lead to an increase in the number of hibernating ribosomes that are protected from ribophagy during cellular stress. This has the potential not only to modulate the proteome of cancer cells, but also to ensure a reserve of ribosomes which could resume translation after an instance of cellular stress such as hypoxia or chemotherapy. Indeed, recent work in the intestine demonstrated that Mex3a-positive stem cells are resistant to chemotherapy^13^.

Our work uncovers previously unappreciated roles for Mex3a in neuronal differentiation, adding to the knowledge of how cells differentiate from stem cells to “outward” facing differentiated cells. We expect that the molecular mechanisms and targets of Mex3a revealed here will provide insight into contexts where Mex3a is dysregulated, as in cancers. These data will also guide therapeutic strategies with the aim of differentiating stem cells to replace diseased cells for patients with neurodegenerative disease.

### Limitations of the study

We observe Arl13b in dendritic knobs of mature OSNs, while a recent study found Arl13b to be restricted to primary cilia of immature OSNs^114^. We suspect that a difference in antibodies may account for this discrepancy. We do not seek to comment on Arl13b protein localization or function, only the ciliary structures illuminated by immunofluorescence. We used SEM to confirm our results in an antibody independent way.

We use HEK293T cells as a model system to explore Mex3a function in addition to the conditional knockout mouse. The cell culture system has the advantage of testing hypotheses rapidly with genetic constructs as well as testing whether findings might be evolutionarily conserved to human, but it has the disadvantage that transfection of exogenous Mex3a could lead to increased cellular stress or dysregulation of homeostatic cellular processes.

Our live cell analysis of vesicles in cultured OSNs is a first step towards exploring vesicle quantities and dynamics in neurons. We rely on the fact that the GFP from the endogenously expressed Ngn1-GFP allele does not enter structures in the cell that are bounded by membranes. We used careful training of NIS software and provided cutoffs related to circularity and size when training Segment.ai in order to ensure that cytoplasmic vesicles would be counted instead of other membrane-bound organelles like Golgi or ER. Further experiments should be conducted by repeating the experiment with labeled markers of cytoplasmic vesicles.

The ubiquitin linkage IP-Western experiment is performed in HEK293T cells. We tested the most likely ubiquitin chain that Mex3a confers on ubiquitin targets by co-transfecting a Mex3a construct with a putative ubiquitin target (Serbp1 or Rps7) and an HA-ubiquitin construct. This experiment demonstrates that Serbp1 and Rps7 can be K27 ubiquitinated, however, E3 ubiquitin ligases expressed in HEK293T cells, including endogenous human MEX3A itself, may contribute to the K27 signal we observe. Further experiments would be necessary to demonstrate definitively that Mex3a confers the K27 linkage.

Our Polysome profile experiments are conducted with mixed populations of cells: for mouse we use whole MOE tissue, and for HEK293T cells, a portion of the cells are transiently transfected, and a portion are not. Due to these mixed populations, our results may underestimate the role of Mex3a in regulating polysome to monosome ratios, recruitment of Serbp1 and p-eEF2, and translational elongation and/or initiation.

## Supporting information

Supplemental File 1 Total Proteome

Supplemental File 2 HyperTRIBE

Supplemental File 3 Ubiquitin Mass Spec Raw

Supplemental File 4 Ubiquitin Mass Spec Candidates

Supplemental File 5 DESeq Polysome Profiling RNA Seq

## Resource Availability

Further information and requests for resources and reagents should be directed to and will be fulfilled by the Lead Contacts, Rachel Duffié, Stavros Lomvardas, Marko Jovanovic

- Data are deposited in GEO (accession numbers GSE300224 and GSE300225) and MassIVE (<u>MSV000100330</u>)
- Plasmids constructed for this study and used to make transgenic HyperTRIBE mice, and for transient transfections in HEK293T cells can be found at Addgene (ID numbers 241312, 241313, 241314, 241315, 241316, 241318). Any additional information required to reanalyze the data reported in this paper is available from the lead contacts upon request.

## Acknowledgements

We thank Drs. Daniele Canzio and Franck Polleux for critical reading of the manuscript. We are grateful to Dr. Alex Baffet, Jovanovic lab members especially Drs. Lena Annika Street and Ella Doron, and Lomvardas lab members for critical discussion. We acknowledge the expertise and support from the animal husbandry teams at Columbia University, the Zuckerman Institute Flow cytometry core, the Flow Cytometry Core of the Columbia Center for Translational Immunology (CCTI) and Herbert Irving Comprehensive Cancer Center (HICCC). Imaging was performed with support from Humberto Ibarra Avila and the Zuckerman Institute’s Cellular Imaging platform. Scanning Electron Microscopy was conducted with the help of Dr. Amir Zangiabadi and facilities and instrumentation supported by NSF through the Columbia University, Columbia Nano Initiative, and the Materials Research Science and Engineering Center DMR-2011738.

## Author Contributions

Conceptualization: RD, SL, and MJ. Formal Analysis: RD, HS, AH, MJ. Funding Acquisition: SL, MJ, RD, LCT. Investigation: MEA, LCT, RD, AK, AU, KA, OS, JY, JP. Methodology: MEA, LCT, AK, AU. Resources: SL, MJ, JY. Visualization: RD. Writing – original draft: RD. Writing – review and editing: MEA, MJ, RD.

## Materials and Methods

### Mice

Mouse protocols were approved by the Columbia University IACUC under protocol numbers AC-AAAT2450 and AC-AABG6553. All mice were housed in standard conditions with a 12-hour light/dark cycle and access to food and water *ad libitum*. Animals were on a mixed genetic background and littermate controls were used for comparisons. Animals were sacrificed by CO_2_ followed by cervical dislocation, and the main olfactory epithelium was isolated by dissection.

Transgenic mice were made with the help of Dr. Nataliya Zabello in Dr. Richard Axel’s laboratory. They were constructed with the following plasmids, tetO-CMV-Mex3a-HyperTRIBE-V5-t2a-mCherry (Addgene ID 241315) and tetO-CMV-HyperTRIBE-V5-t2a-mCherry (Addgene ID 241314). Mouse Adar deaminase domain was added by In Fusion cloning (Takara) of gene block (IDT) to pTRE2-CMV-Mex3a-t2a-mCherry (Addgene plasmid 223244) digested with HindIII (NEB). To generate the tetO-CMV-HyperTRIBE-V5-t2a plasmid, the tetO-Mex3a-HyperTRIBE plasmid was digested with NheI and ClaI to remove the Mex3a gene and repaired by ligation of annealed oligos with overhangs.

| **Mouse Line** | **Publication** | **Source/Identification** |
| --- | --- | --- |
| *Mex3a flox* | Álvarez-Varela et al 2022 | Donated by PI |
| *Foxg1tm1.1(cre)Ddmo/J* | Kawaguchi et al 2016 | Jax #029690 |
| *Ngn-GFP, Tg(Neurog1-EGFP)DF148Gsat* | MMRRC, Gong et al 2003 | #010618-UCD |
| *Omp(iresGFP)* | Shykind et al 2004 | Donated by PI |
| *tetO-HyperTRIBE-V5-t2A-mcherry* | This study | Generated in house |
| *tetO-Mex3a-HyperTRIBE-V5-t2a-mCherry* | This study | Generated in house |
| *gg8-tTA* | Nguyen et al 2007 | Donated by PI |
| *Omp(irestTA), Omptm1(tTA)Gogo "OMP-tTA"* | Yu, C.R. et al 2004 | Jax #017754 |

### Cell Culture

HEK293T cell line was obtained from ATCC, grown at 37°C; 5% CO2 in a humidified incubator and regularly screened for mycoplasma. HEK293T cells were maintained in high glucose Dulbecco’s modified Eagle’s medium (DMEM) (Gibco) supplemented with 10% heat inactivated Fetal Bovine Serum (Corning), 1X Minimum Essential Medium Non-essential amino acids (Gibco) and 1X GlutaMAX (Gibco). HEK293T cells were grown without antibiotics.

### Immunofluorescence

MOE was dissected and frozen in OCT compound (Fisher) and sectioned at 12 µM with a cryostat (Leica) onto a microscope slide (Fisher Scientific). Tissue sections were dried at RT for ten minutes, fixed with 4% PFA, 1X phosphate-buffered saline (PBS) for 5 minutes, washed with 1X PBS, then blocked for 30-60 minutes in blocking solution (4% Sterile Donkey Serum (Sigma), 1% Triton X-100, 1X PBS). Slides were incubated in a humid chamber overnight with primary antibody diluted 1:200 in blocking solution, with a coverslip placed on top to avoid evaporation. The following day, coverslips were removed by adding 1X PBS to the top, slides were washed in a slide mailer 3×5 minutes in PBST (1X PBS, 0.1% Triton X-100), then incubated in blocking solution with secondary antibody (diluted 1:500) and DAPI (diluted 1:1000) before mounting with Vectashield mounting media (Vector Labs). Images were acquired using a a Zeiss LSM 700 Confocal, a W1-Yokogawa spinning disk Confocal, or a Zeiss LSM 800 series Confocal microscope with Airyscan, depending on experiment.

For immunofluorescence images where endogenous GFP is visualized (Ngn1-GFP or OMP-ires-GFP), MOE was dissected, fixed for 8 minutes on ice in 4% PFA; 1X PBS, washed three times with 1X PBS, and incubated overnight in 30% sucrose; 1X PBS. Tissue was mounted in OCT and IF experiment continued as described above.

Immunofluorescence analysis: All analyses for images acquired after immunofluorescence were completed in FIJI and R studio. Average OR Signal Intensity: FIJI freehand manual selection tool was used to manually delineate each OR positive cell and assign it to a region of interest (ROI). Fluorescence intensity was then measured for each cell in the ROI list. Delineating OR positive cells into apical or basal segments of the tissue was described in^8^. Radial profile of Arl13b: FIJI circle selection tool was used to circle each dendritic knob, then radial profile of Arl13b signal was quantified using FIJI and plotted with R studio. Average Adcy3 signal intensity: Entire coronal sections of the MOE were imaged, and 160 µM x 160 µM sections were selected across the epithelium to ensure all zones were represented. Mean Fluorescence Intensity was quantified for 32 tiles from each genotype. Polarity analysis: To quantify polarity, a line was drawn with the line selection tool in FIJI directly through the Tgn46 signal for each OR positive cell, and the angle was measured between this line and a line along the apical to basal axis of the tissue, using the FIJI angle tool.

Antibodies used for immunofluorescence:

| Antibody | Source |
| --- | --- |
| Rabbit anti Mex3a | Abcam 79046 |
| Goat anti Calmegin N-16 | Santa Cruz 49899 |
| Rabbit/Guinea Pig anti P2, C7, M71, Mor28 OR proteins | Generous gift from Gilead Barnea, Brown University |
| Rabbit anti Arl13b | Abcam 83879 |
| Rabbit anti Tgn46 | Abcam 16059 |
| Rabbit anti Adcy3 | Santa Cruz 588 |

### Scanning Electron Microscopy

MOE was carefully dissected, and surrounding bone tissue was meticulously removed. The sample was fixed in 2% glutaraldehyde in 1X PBS for one hour at 4°C, followed by three 10-minute washes in 1X PBS. Dehydration was performed in a graded ethanol series: 25% ethanol for 15 minutes, 50% ethanol overnight, 75% ethanol for four hours the following day, and 100% ethanol overnight.

The sample was then placed inside a microporous specimen capsule and dried using a critical point dryer (CPD). Once completely dry, the specimen was mounted on an aluminum stub for scanning electron microscope (SEM) and coated with a 60:40 gold/palladium alloy. Imaging was performed using a Zeiss Sigma VP SEM with an accelerating voltage of 12 kV to ensure high image quality and stability.

SEM analysis: To quantify cilium position within the dendritic knob, FIJI point tool was used to generate a ROI list with each site of origin for each cilium projection, as well as a ROI delineating the extent of the dendritic knob (circle area selection tool) for each visible knob in our images. R studio was used to set the radius of each dendritic knob to 1, and the origin points were normalized to this adjusted size and plotted. Their distance from the center of the knob was measured, and statistical significance between genotypes was calculated using the Wilcoxon rank sum test.

### Transient Transfection of HEK293T cells

Transient transfection was performed with Lipfectamine 3000 reagent. 700K Cells were seeded (Day 0) on Gelatin coated (EmbroMax 0.1% Gelatin Solution (Merck Millipore)) 6-well plates and transfected with a total of 2.5µg of DNA the following day (Day 1). If two plasmids were transfected, 1.25µg of each plasmid was used, if three plasmids were transfected, 0.83µg of each plasmid was transfected. Cells were passed 1:4 the following day (Day 2) for imaging or IP, or two identically transfected six-well plates were passed into a 10cm plate for polysome experiments. Experiments were started on Day 3, two days after transfection.

Plasmids used for transient transfection of HEK293T cells:

| **Plasmid** | **Source** |
| --- | --- |
| CMV-Mex3a-mCherry | This study, available at Addgene 241312 |
| CMV-mCherry | This study, available at Addgene 241313 |
| pcDNA3/mStayGold(c4)=UtrCH | Addgene 212020 |
| GFP-Sec61b | Addgene 121159 |
| CMV-Serbp1-mEGFP | This study, available at Addgene 241316 |
| CMV-Rps7-mEGFP | This study, available at Addgene 241318 |
| CMV-mEGFP | This study, Addgene 129590 cut with EcoRI, and repaired to add a start codon |
| pRK5-HA-ubiquitin WT construct | Addgene 17608 |
| pRK5-HA-ubiquitin K0 construct | Generous gift from Junqiang Ye |
| pRK5-HA-ubiquitin K6 construct | Addgene 22900 |
| pRK5-HA-ubiquitin K11 construct | Addgene 22901 |
| pRK5-HA-ubiquitin K27 construct | Addgene 22902 |
| pRK5-HA-ubiquitin K29 construct | Addgene 22903 |
| pRK5-HA-ubiquitin K33 construct | Addgene 17607 |
| pRK5-HA-ubiquitin K48 construct | Addgene 17605 |
| pRK5-HA-ubiquitin K63 construct | Addgene 17606 |

### Live Cell Imaging of HEK293T cells

After transient transfection with CMV-Mex3a-mCherry, CMV-mCherry, pcDNA3/mStayGold(c4)=UtrCH, and/or GFP-Sec61b^29^ plasmids as described above, HEK293T cells were passed into a 35mm glass bottom dish (MatTek) coated with Gelatin (EmbryoMax 0.1% Gelatin Solution) and imaged using a SoRA-W1 Yokogawa Spinning Disk Confocal with 5% CO2 and at 37°C. For each image, 51 slices were taken in Z with a step-size of 0.1 µM. Images were deconvolved using the FIJI plugin Microvolution.

### Culturing and Live Cell Imaging of Mouse olfactory sensory neurons

Performed as described in^117^, briefly, MOE was dissected from early postnatal (PN3) Mex fl/fl; Foxg1iresCre; Ngn1-GFP mice and their Foxg1iresCre negative littermates (1 male and 1 female of each genotype for 2 biological replicates per genotype). Single cell suspension was generated using Worthington Papain dissociation kit (Worthington Biochemical) according to manufacturer’s instructions. Cells were filtered using a 40 µM Flowmi cell strainer, pelleted, and resuspended in mOSN medium (Waymouth’s MB 752/1 medium (Gibco, 11220035), 1% N2 supplement (100X Gibco, 17502048), 1% Antibiotic-Antimycotic (100X, Gibco, 15240062)). 600,000 cells were plated per 35mm^2^ dish of confluent astrocytes. OSN cultures were maintained at 37°C with 5% CO_2_ in mOSN medium. 24 hours after plating, half of mOSN medium was replaced with mOSN medium and from then, half of medium was replaced every other day. Imaging was conducted on Day 3 post dissection/plating using a SoRA-W1 Yokogawa Spinning Disk Confocal with 5% CO2 and at 37°C. For each image, 51 slices were taken in Z with a step-size of 0.1 µM. Images were deconvolved using the FIJI plugin Microvolution. Vesicles were identified using NIS software by training the Segment.AI function, with additional filters for size and circularity applied. WT and cKO images were batched processed with the same segmentation algorithm to ensure identical analysis pipeline.

### Fluorescence Activated Cell sorting

MOE was dissected and a single cell suspension was generated using the Worthington Papain dissociation system according to manufacturer’s protocol (Worthington Biochemical). After trituration and inactivation of papain, cells were resuspended in sort medium (2% FBS; 1X PBS for cells used in Mass Spectrometry experiments to reduce contaminating peptides, 10% FBS; 1X PBS for cells used in RNA-Seq experiments) and filtered with a 40 µM Flowmi cell strainer (Fisher Scientific). Cells were sorted on a Beckman Coulter MoFlo Astrios EQ Cell Sorter or a BD Influx Cell Sorter and centrifuged at 500 rpm in a swinging bucket rotor to remove sort medium. Cells for Mass Spectrometry were washed three times with 1X PBS to remove FBS. Cells for RNA-Seq were resuspended in 500 µL TRIzol Reagent (Invitrogen).

### Total proteome mass spectrometry

#### Sample Preparation

Sorted cell pellet was resuspended in Urea lysis buffer (8M urea, 75mM NaCl, 50mM Tris pH8, 1mM EDTA, 1X HALT protease inhibitor (Thermo Fisher Scientific) at a ratio of 130 µL per million cells. Proteins were reduced by adding 4mM DTT and incubating 45 minutes at RT, shaking 1,000 rpm. Proteins were alkylated by adding 10mM iodoacetamide, protected from light and incubated 45 minutes RT, shaking 1,000 rpm. Peptides were digested by adding 0.5ug Trypsin and incubating at RT overnight, shaking 700 rpm. The following day, 1% formic acid was added to acidify proteins. C18 stage tips were prepared by packing two disks of Empore 3M C18 material into 200uL tips. Stage tips were equilibrated by sequential washes of 100 μL of 100%MeOH, 80%ACN/0.2%Formic Acid, 2x 3%ACN/0.2%Formic Acid, then the acidified peptides were loaded on the stage tip. Stage tips were washed twice with 3%ACN/0.2% Formic acid, and peptides were eluted with 60% ACN/0.2% Formic Acid, dried in a SpeedVac vacuum concentrator.

#### LC-MS/MS Parameters

Whole proteome, label-free MS analysis was performed by data-independent acquisition (DIA). For this type of LC-MS/MS analysis, about 1 μg of total peptides were analyzed on a Waters M-Class UPLC using a 15cm IonOpticks Aurora Elite column (75mm inner diameter; 1.7mm particle size; heated to 45°C) coupled to a benchtop Thermo Fisher Scientific Orbitrap Q Exactive HF mass spectrometer. Peptides were separated at a flow rate of 400 nL/min with a 100 min gradient, including sample loading and column equilibration times. Data was acquired in data independent mode. MS1 Spectra were measured with a resolution of 120,000, an AGC target of 3e6 and a mass range from 350 to 1650 m/z. Per MS1, 34 equally distanced, sequential segments were triggered at a resolution of 30,000, an AGC target of 3e6, a segment width of 38 m/z, and a fixed first mass of 200 m/z. The stepped collision energies were set to 22.5, 25, and 27.

#### Analysis

All DIA data were analyzed with Spectronaut software version 15.7.220308.50606 (Bruderer R et al 2015) using directDIA analysis methodology against a Mus musculus Uniprot database. Carbamidomethylation on cysteines was set as a fixed modification. Oxidation of methionine and protein N-terminal acetylation were set as variable modifications. Trypsin/P was set as the digestion enzymes. Raw protein group quantities were normalized by library depth (raw quantity/(sum of all raw quantities measured for that sample * 1,000,000). Log2 values for each protein group was averaged across replicates, and LFC was calculated by subtracting (Mex3a cKO – WT). To generate high confidence lists presented in Figure 2D and E, a Student’s t-test was performed to test significance of LFC Mex3a cKO vs WT. Proteins were considered statistically significant if 1) they had Student’s t-test value of less than 0.05, 2) log fold changes (Mex3a cKO – WT) of greater than 0.58 or less than -0.58, and 3) were measured in all 7 samples to ensure sufficient coverage of a given protein.

### *In vivo* HyperTRIBE in mouse

To test TRIBE constructs, the deaminase domains of mouse Adar, Adarb1, and Adarb2 were cloned in frame with an MS2 coat protein and recruited to GFP RNA using a GFP-Ms2-stem loop construct by co-transfecting both constructs into HEK293T cells. RNA was extracted with TRIzol, DNase I treated (DNase I, Ambion) followed by cDNA conversion (SuperScript III, Invitrogen). GFP cDNA was amplified by PCR with Herculase II polymerase (Agilent) to ensure low error rate, and reactions were incubated with GoTAQ polymerase to add A-overhangs. Products were Topo cloned (Invitrogen) and mini preps were Sanger sequenced to identify editing events. This assay revealed that the Adar deaminase domain led to the highest level of editing (data not shown). The mouse Adar deaminase domain with the HyperTRIBE mutation was cloned downstream of the tetO-Mex3a construct previously used to make transgenic mice, which harbors an mCherry to facilitate FACs^8^. See “Mice” section in Materials and Methods for additional cloning details.

Three transgenic founders of tetO-HyperTRIBE only and six transgenic founders of tetO-Mex3a-HyperTRIBE were obtained, and all founders were bred gg8-tTA; OMP-tTA mice. Progeny were tested for expression of the transgenes by immunofluorescence for the V5 tag in the constructs (data not shown). After all constructs were found to exhibit widespread expression across the MOE, additional progeny were used for RNA-Seq experiments. MOE from progeny (age 3-5 weeks) of each founder was dissected, and individual cells were isolated using Worthington Papain and FAC sorted as described above into mCherry positive and negative cell populations. RNA was extracted from mCherry positive and negative samples as described above, and 10ng was used as input for RNA-Seq libraries, which were prepared as described below.

Bioinformatics was performed with tools described in^30^, with modifications to account for CIGAR score differences in mouse. RNA-Seq from mCherry negative cells was used as a negative control for comparison to mCherry positive cells from the same mouse, which accounts for SNP genetic differences from mouse to mouse. Outputs from each mouse were compared to biological replicates. For each RNA, a HyperTRIBE score was generated based on how many editing events were measured for that RNA and how many biological replicates exhibited editing for that RNA.

To identify putative Mex3a RNA targets, two tetO-Mex3a-HyperTRIBE founders were bred to gg8-tTA mice to express the transgene in immature OSNs where Mex3a is expressed endogenously. FAC sorting, RNA-Seq, and bioinformatics were performed as described above and the top 15% RNAs with the highest HyperTRIBE scores were considered putative Mex3a targets.

### RNA-Sequencing

Sorted cells were lysed at RT in TRIzol for 10 minutes, then 1/5 Trizol Volume (100 µL) of 1-Bromo-3-chloropropane (Sigma Aldrich) was added to isolate RNA into the upper aqueous phase. Isopropanol was used to precipitate RNA with 1 µL GenElute LPA (Sigma Aldrich) to facilitate precipitation. The RNA pellet was washed with 75% Ethanol, allowed to dry 5 minutes at RT, and resuspended in H_2_O. RNA was immediately treated with DNase I (Ambion) at 37°C for 30 minutes, and purified using AMPure XP beads (Beckman Coulter). RNA-Seq libraries were made using the SMARTer Stranded Total RNA-Seq Kit - Pico Input (Takara) according to manufacturer’s protocol using 10ng of RNA as input. Library was quantified using the KAPA library quantification kit (Roche) and quality was confirmed using an Agilent Bioanalyzer before sequencing on a NextSeq 550 Illumina machine.

RNA was extracted from polysome samples using TRIzol LS reagent. 50 µL each from four fractions were pooled (200 µL Sample + 600 µL TRIzol LS). After lysis, 160 µL of 1-Bromo-3-chloropropane (Sigma Aldrich) was added and aqueous upper phase was combined for Early or Late polysomes (Fractions 27-34 are Early Polysomes, Fractions 35-42 are Late Polysomes). 320µL of ETOH 100% was added and the samples were purified using the Zymo RNA Clean & Concentrator kit (Zymo) to eliminate sucrose, followed by elution in 15 µL of H_2_O. 10ng RNA were used as input for the SMARTer Stranded Total RNA-Seq Kit – Pico Input (Takara). Library was quantified with Qubit (Thermo Fisher Scientific) and quality was confirmed with an Agilent Tapestation and sequenced on a HiSeq 2500 Illumina machine.

RNA-Seq data were aligned to the mm10 genome with STAR and analyzed using DESeq2 in R.

### UbiFast Mass Spectrometry

Six Mex3a cKO and six WT littermate controls were collected (age PN12, two MOEs pooled for three biological replicates each genotype) and snap frozen in liquid nitrogen. Tissue was cryosmashed using a Covaris CP02 tissue pulverizer and lysed in 1mL of Urea lysis buffer (see above) for 20 minutes at RT rotating end over end. Samples were centrifuged at 14,000 g for 10min at 12°C to remove cell debris. Supernatant was collected and quantified using the Pierce BCA Protein Assay (Thermo scientific). 2mg of protein was used for each IP.

Before immunupurification, proteins were reduced by adding 4mM DTT and incubating 45 minutes at RT, shaking 1,000 rpm. Proteins were alkylated by adding 10mM iodoacetamide, protected from light and incubated 45 minutes RT, shaking 1,000 rpm. Peptides were digested by adding 0.5µg Trypsin and incubating at RT overnight, shaking 700 rpm. The following day, 1% formic acid was added to acidify peptides. C18 stage tips were prepared by packing two disks of Empore 3M C18 material into 200µL tips. Stage tips were equilibrated by sequential washes of 100 μL of 100%MeOH, 80%ACN/0.2%formic acid, 2x 3%ACN/0.2%formic acid, then the acidified peptides were loaded on the stage tip. Stage tips were washed twice with 3% ACN/0.2% formic acid, peptides were eluted with 60% ACN/0.2% formic acid, and dried in a SpeedVac vacuum concentrator.

Peptide samples were brought up to 1.5mL with 1X HS IAP Bind buffer (from Cell Signaling Technology PTMScan HS Ubiquitin Remnant Motif (K-ε-GG) Kit #59322) and sonicated with a Branson 2800 cleaner on “High” setting for 5-10 minutes to resuspend peptides. pH was confirmed at 7 or adjusted with 1M Tris. Peptides were spun down at 10,000g for 5 minutes and supernatant was kept. 20 µL Magnetic K-ε-GG beads per IP were washed 4X with ice cold 1X PBS then resuspended in IAP Bind buffer. Reconstituted peptides were added to K-ε-GG bead aliquots and incubated overnight at 4°C, rotating end over end. The following day, washes were performed with chilled solvents on ice as quickly as possible. Immunopurified samples were collected by magnet and flow-through was removed. Beads were washed 4X with 1mL 1X HS IAP Wash buffer. Next, beads were washed 2X with 1mL HPLC grade water. K-ε-GG peptides were eluted with two 10-minute elution steps at RT with 50 µL 0.15% Trifluoroacetic acid, gently flicking beads into solution every 2-3 minutes. Supernatant was collected then concentrated and purified using C18 stage tips. C18 stage tips were prepared by packing two disks of Empore 3M C18 material into 200 µL tips. Stage tips were equilibrated by sequential washes of 100 μL of 100%MeOH, 80%ACN/0.2%formic acid, and 2x 1% formic acid, then the immunopurified peptides were loaded on the stage tip. Stage tips were washed twice with 1% formic acid, peptides were eluted with 60% ACN/0.2% formic acid, and dried in a SpeedVac vacuum concentrator.

#### LC-MS/MS Parameters were as described above

Ubifast data were quantified by Spectronaut with standard settings as described above, using the ubiquitin post-translational modification search. The data were analyzed both with global imputation and without imputation. Protein candidates presented in Figure 4 were statistically significant (Q-value less than 0.05) in either or both of the analyses.

### Ubiquitin Linkage Assay

HEK293T cells were seeded into 6 well plates and transiently transfected (Day 0) as described above with 0.83µg each of Mex3a-mCherry and Serbp1-mEGFP, Rps7-mEGFP, or mEGFP and one of 9 pRK5-HA-ubiquitin constructs. Initial tests showed that K48 input samples showed very little HA staining by Western blot, so all cells were treated with a final concentration of 20 µM MG132 (Sigma) on Day 1. MG132 treatment did not affect abundance of other ubiquitin constructs as determined by Western blot (data not shown). On Day 2 after transfection, cells were trypsinized, centrifuged with a swinging bucket rotor and lysed in 400 µL Lysis buffer (150mM NaCl, 50mM Tris pH7.5, 1% NP-40, 5% Glycerol, 1X HALT protease inhibitors, and 1X phosphatase inhibitors). Cells were lysed for 30 minutes at 4°C rotating end over end, then centrifuged at max speed for 10 minutes at 4°C. 20 µL of lysis-buffer washed Anti-GFP mAb-Magnetic beads (MBL international) were added to supernatants, and immunopurification proceeded overnight at 4°C. The following day, beads were washed three times with Wash buffer (150mM NaCl, 50mM Tris pH7.5, 5% Glycerol, 0.05% NP-40). Protein was eluted off beads at 70°C for 10 minutes in SDS Western loading buffer (30 µL Lysis buffer + 10 µL 4X Laemmli with β-mercaptoethanol), and Western blot was performed as described below.

### Polysome Profiling

Mouse polysome fractionation: MOE was dissected and washed with ice cold 1X PBS supplemented with 100µg/mL Cyclohexamide. Tissue was flash frozen in liquid nitrogen and stored at -80°C until day of polysome fractionation experiment. Each sample consisted of ten MOEs PN12 Mex3a cKO and their Mex3a WT littermates, two biological replicates (20 MOEs total per genotype). On day of polysome fractionation, sucrose gradients were prepared as follows: 10X polysome gradient buffer (206mM Hepes-KOH pH 7.4, 20.6mM Mg(OAc)_2_, 1M KOAc) was taken to 1X in 10% and 50% sucrose solutions. Gradients were made in ultracentrifuge tubes (Beckman Coulter 344059). 10% sucrose solution; 1X polysome gradient buffer was added to the tube, and 50% sucrose solution; 1X polysome gradient buffer was carefully layered underneath with a syringe. Tubes were capped and gradient was made with a gradient maker (BioComp instruments), then chilled at 4°C for 30min-1hour.

During this time, MOE samples were cryosmashed in a Covaris CP02 tissue pulverizer, and lysed with polysome lysis buffer (10mM Tris-HCl pH 7.5, 100mM KCl, 5mM MgCl_2_, 0.5% sodium deoxycholate, 1% NP-40, 10mM DTT, 100µg/mL Cyclohexamide, 1U/mL SUPERase-In, and 1X Protease inhibitor) was added (1mL lysis buffer for 10 MOEs). Tissue was pipetted repeatedly to lyse completely, allowed to sit on ice two minutes, pipetted again, then transferred to a new tube and incubated an additional three minutes on ice. Samples were centrifuged in a tabletop centrifuge 2000g for five minutes at 4°C, supernatant was transferred to a new tube and centrifuge step was repeated. 400 µL MOE lysis was added onto chilled gradients then ultracentrifuged (Beckman Coulter XPN-100, SW41TI rotor) at 38,000 rpm for 2 hours at 4°C. A BioComp Fractionater was used to measure absorbance and collect ∼42 fractions per condition.

HEK293T polysome profiling: HEK293T cells were seeded and transiently transfected in 6 well plates as described above with mCherry + Serbp1-mEGFP or Mex3a-mCherry + Serbp1-mEGFP plasmids, ten wells for each condition. The day after transfection, cells from 2 six-well plates were passed into a 10cm (five 10cm per condition) and allowed to grow an additional day. Sucrose gradients were prepared as described above. During that time, cell lysates were prepared as follows: Cyclohexamide was added to medium at a final concentration of 100µg/mL, swirled, and incubated at 37°C for 1-2 minutes. Plates were placed on ice and cells were washed two times with 5mL ice cold PBS 1X supplemented with 100µg/mL Cyclohexamide. Wash was aspirated carefully and completely and 400 µL Lysis buffer (Polysome buffer: 20mM Tris pH 7.56, 150mM NaCl, 5mM MgCl_2_, 100µg/mL Cyclohexamide, 1mM DTT, 8% glycerol. Lysis buffer: Polysome buffer supplemented with 1% Triton X-100, 12 U/mL Turbo DNase, 1.6 µL/mL SUPERase-In (Thermo)) was added to first dish of each condition. Cells were scraped with lysis buffer and this suspension was moved to the next plate of that condition until all 5 plates had been lysed. Lysate was transferred to a 1.5mL Eppendorf tube and pipetted 15 times to ensure complete lysis. Samples were centrifuged in a tabletop centrifuge at max speed (20,000g) for 10 minutes at 4°C. Supernatant was collected to a new tube and loaded onto a chilled sucrose gradient, then ultracentrifuged as described above.

Polysome profiles were displayed and quantified with QuapPro^116^. Area under monosomes and polysomes were quantified and used for further analysis.

### Western blot

Loading buffer: 4X Laemmli buffer with 10% β-mercaptoethanol was added to each sample to a final concentration of 1X loading buffer, and samples were incubated at 70°C for 10 minutes. Samples were loaded on a 4-20% mini-PROTEAN TGX precast protein gel (Bio-Rad), ladder: Precision Plus protein dual color standards (Bio-Rad) and subjected to SDS-PAGE. After electrophoresis, proteins were transferred to a PVDF membrane for 90 minutes at 250mA constant. Membranes were blocked in Odysee Blocking buffer (LiCOR) before incubating overnight with primary antibodies diluted 1:3000 in blocking buffer, supplemented with 0.2% Tween-20. The following day, membranes were washed 3×7 minutes with 1X PBS; 0.1% Tween-20, then incubated with LiCOR secondary antibodies (diluted 1:10,000 in LiCOR blocking buffer supplemented with 0.2% Tween-20 and 0.01% SDS) for 1 hour at RT. Membranes were washed 3×7 minutes with 1X PBS; 0.1% Tween-20 and imaged using a LiCOR Odysee F Imaging system. Western band intensity was quantified using LiCOR software. For polysome profiling Westerns, membranes were first blotted for Serbp1 and Rpl30, then after imaging and quantifying bands, the membrane was washed 3×7 minutes with 1X PBS; 0.1% Tween-20 before incubating overnight with p-eEF2 antibody diluted 1:3000 in Odysee Blocking buffer. The following day, washes and secondary antibody incubation was carried out as described above, and the blot was re-imaged on a LiCOR Odysee F Imaging system, quantifying p-eEF2 bands.

Antibodies used for Western blot:

| Antibody | Source |
| --- | --- |
| Rabbit anti HA | Abcam 9110 |
| Chicken anti GFP | Abcam 13970 |
| Mouse anti Serbp1 | Thermo Fisher MA5-26478 |
| Rabbit anti p-eEF2 (thr56) | Thermo Fisher 702808 |
| Rabbit anti Rpl30 | Abcam 170930 |
| Rabbit anti Mex3a | Abcam 79046 |

### Quantification and statistical analysis

Imaging experiments were quantified with FIJI or NIS Elements and R. RNA-Seq data were quantified with STAR and DESeq2. Mass Spectrometry data were quantified with Spectronaut and visualized with Morpheus (Broad Institute) and STRING (string-db.org). HyperTRIBE editing was quantified as described in^30^, with additional processing conducted in R. Polysome profiles and Westerns were quantified by BioComp Fractionater, QuapPro, and LiCOR analysis.

Statistical tests were performed in R and specific tests and n values are listed in figure legends and Materials and Methods. A p-value cutoff of 0.05 was considered statistically significant for all analyses. P-values are presented on figures, with asterisks representing confidence as follows: P<0.05 = *, P<0.01=**, P<0.001=***.

---

**Figure S1:**
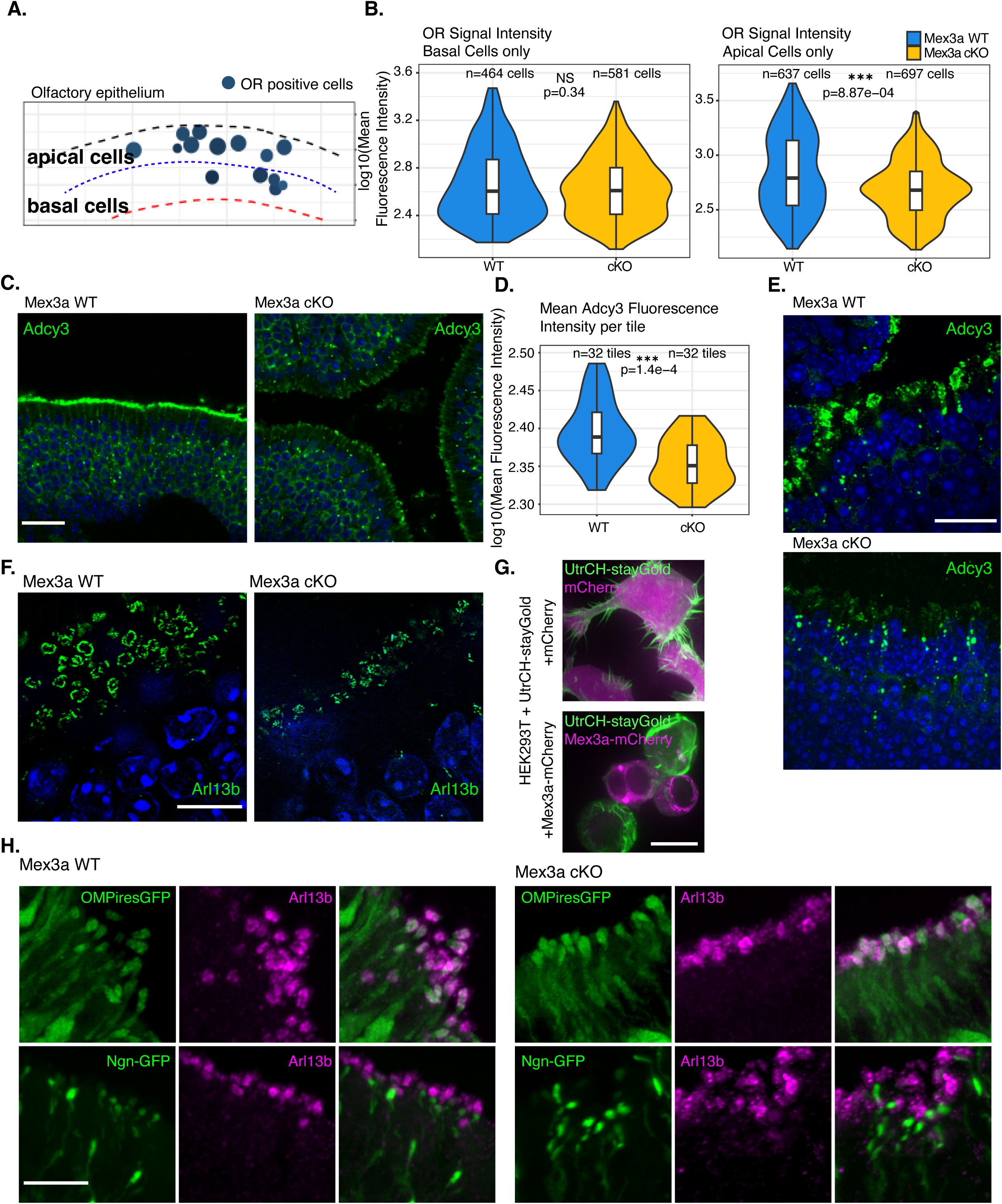
Supporting Main Figure1. **A.** Example representation of image analysis of OR positive cells’ localization in the MOE, same data as in Figure 1B and C. For each image, the apical and basal extent of the MOE was delineated in FIJI, then cells were assigned a position and partitioned into either the basal lower half (more immature/progenitor cell) or apical upper half (more mature OSN) of the epithelium. **B.** Violin plots quantifying OR mean fluorescence intensity per cell, for basal cells (left panels) or apical cells (right panels). **C.** Representative immunofluorescence image for Adcy3 (green), DAPI in blue, from greater than three biological replicates at 4-6 weeks of age. Scale bar, 30 µM. **D.** Violin plot quantifying fluorescence intensity of Adcy3 across MOE of three biological replicates, 4 weeks old. Entire section of MOE was scanned with W1-Yokogawa spinning disk confocal and 32 tiles of 160 µM x 160 µM were saved as regions of interest in FIJI. Mean fluorescence intensity in the green Adcy3 channel was calculated for each tile. Statistics, Wilcoxon rank sum test. **E.** Immunofluorescence for Adcy3, 100X magnification of staining in cilia. Images obtained with W1-Yokogawa spinning disk confocal. Representative image from two biological images at PN12. Scalebar, 20 µM. **F.** Immunofluorescence of Arl13b imaged with Zeiss LSM 800 confocal Airyscan microscope depicting dendritic knobs in apical, airway space above sustentacular and OSN nuclei counterstained with DAPI (blue). Scale bar, 10 µM. **G.** Fluorescence image of UtrCH-stayGold (green) and mCherry or Mex3a mCherry (magenta) in transiently transfected HEK293T cells. UtrCH is an actin binding protein described to mark filapodia and/or lamellipodia. Images taken with SoRA-W1 Yokogawa spinning disk confocal microscope. Scale bar, 10 µM. **H.** Endogenously labeled OMP-ires-GFP (top panels) or Ngn1-GFP (bottom panels) in green with Arl13b immunofluorescence in magenta. Mex3a WT on left and Mex3a cKO on right. Images taken with W1-Yokogawa spinning disk confocal microscope. Scale bar, 10 µM. For all statistical tests in this figure: P<0.05 = *, P<0.01=**, P<0.001=***.

**Figure S2:**
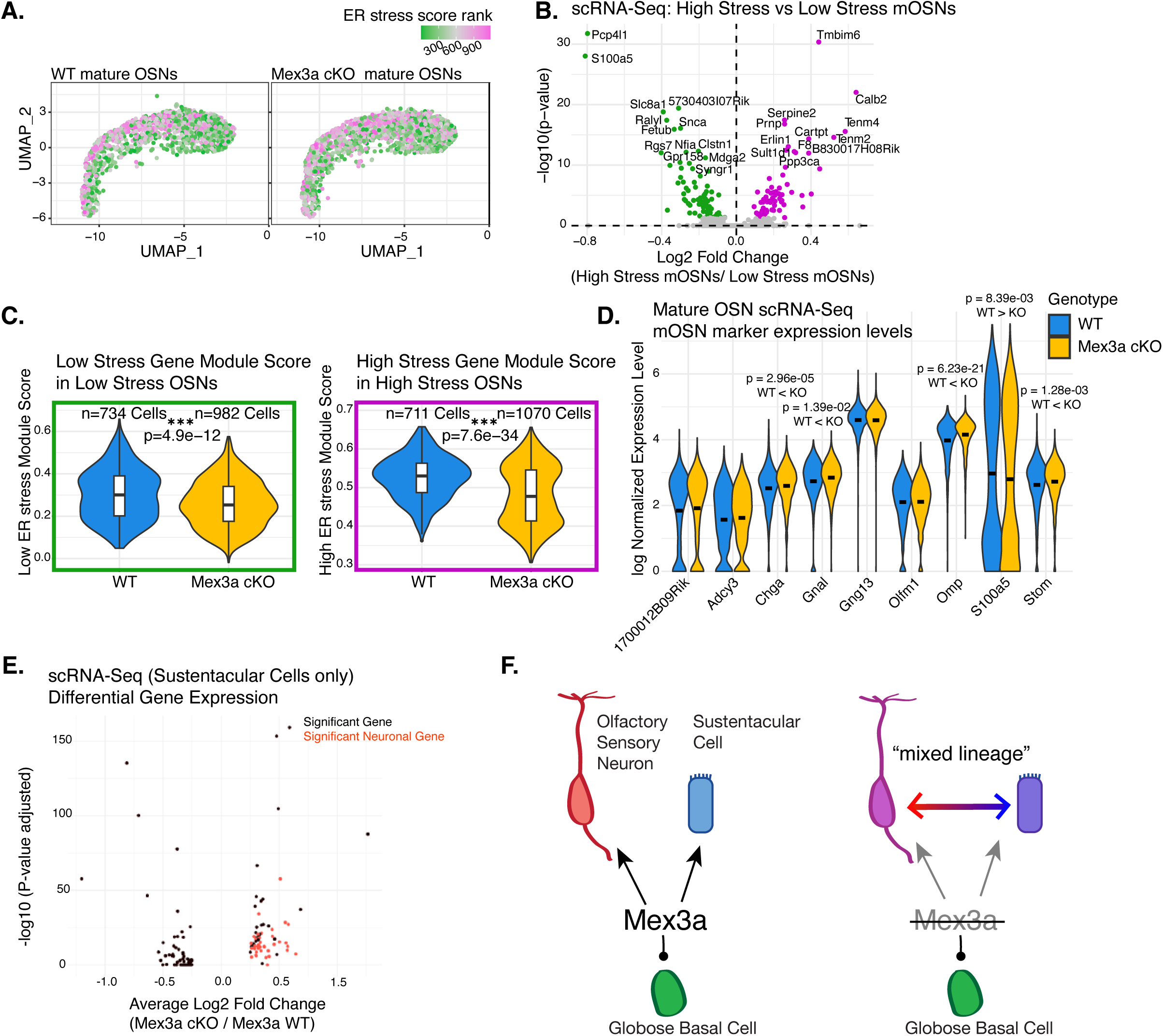
Supporting Main Figure 2. **A.** UMAP projection of mature OSNs in scRNA-Seq experiment described in^8^, Mex3a WT on left, cKO on right. Cells are color-coded by their ER stress rank, which is determined by the chosen OR in each cell^25^ **B.** Volcano plot of differentially expressed genes at RNA level in high and low ER stress Mex3a WT mOSNs in the dataset presented in (A). Differentially expressed genes were used to calculate the high and low ER stress gene module scores quantified in (C). **C.** Violin plots quantifying the Module scores^26^ for “Low ER stress genes” in Low stress mOSNs on the left and “High ER stress genes” in High stress mOSNs on the right for both Mex3a WT and cKO mOSNs. Statistics, Wilcoxon rank sum test. **D.** Log normalized RNA expression levels for known markers of mOSNs in Mex3a WT and cKO mOSNs by scRNA-Seq. Seurat FindMarkers call, Wilcoxon rank sum test. **E.** Volcano plot showing differentially expressed genes (Mex3a cKO sustentacular cells / Mex3a WT sustentacular cells) found by scRNA-Seq of whole MOE, PN12^8^. Genes found to be markers of mature OSNs (Seurat FindMarkers) are colored in red. **F.** Model depicting how Globose basal stem cells, which express high levels of Mex3a, differentiate into olfactory sensory neurons and sustentacular cells. In the absence of Mex3a, we observe changes in differentiation profiles where neurons exhibit higher levels of sustentacular markers, and sustentacular exhibit higher levels of neuronal markers. For all statistical tests in this figure: P<0.05 = *, P<0.01=**, P<0.001=***.

**Figure S3:**
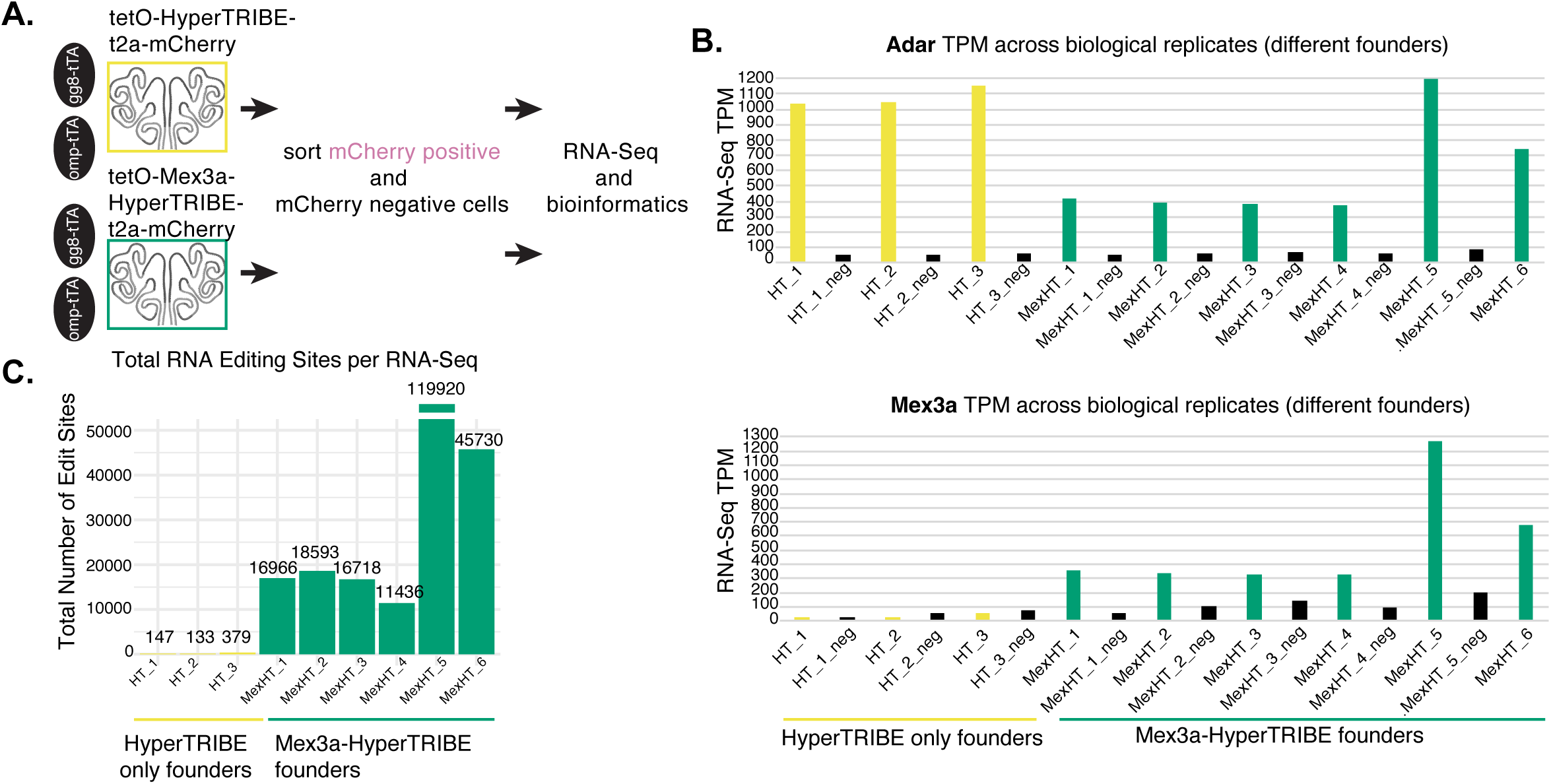
Supporting Main Figure 3. **A.** Experimental design for HyperTRIBE experiment. Negative control (yellow) tetO-HyperTRIBE-V5-t2a-mCherry does not express Mex3a, tetO-Mex3a-HyperTRIBE-V5-t2a-mCherry (green) does express Mex3a. To test the constructs, two MOE-specific drivers, OMP-tTA and gg8-tTA were used, which drive transgene expression in mature and immature neurons, respectively. For putative Mex3a targets, only the gg8-tTA driver was used, as Mex3a is expressed in immature neurons in normal development. **B.** Bar graph plotting transcript per million (TPM) of Adar gene (top) and Mex3a gene (bottom) from RNA-Seq libraries from mCherry+ (colored) or mCherry-(black) sorted cells from individual mice (three different founders for tetO-HyperTRIBE control (abbreviated HT_1 through _3) and six different founders for tetO-Mex3a-HyperTRIBE (abbreviated MexHT_1 through _6)). **C.** Bar graph depicting total number of editing sites (A→I, read out as G after sequencing) per RNA-Seq library of mCherry+ sorted cells. HyperTRIBE only transgenic founders in yellow, Mex3a-HyperTRIBE transgenic founders, green.

**Figure S4:**
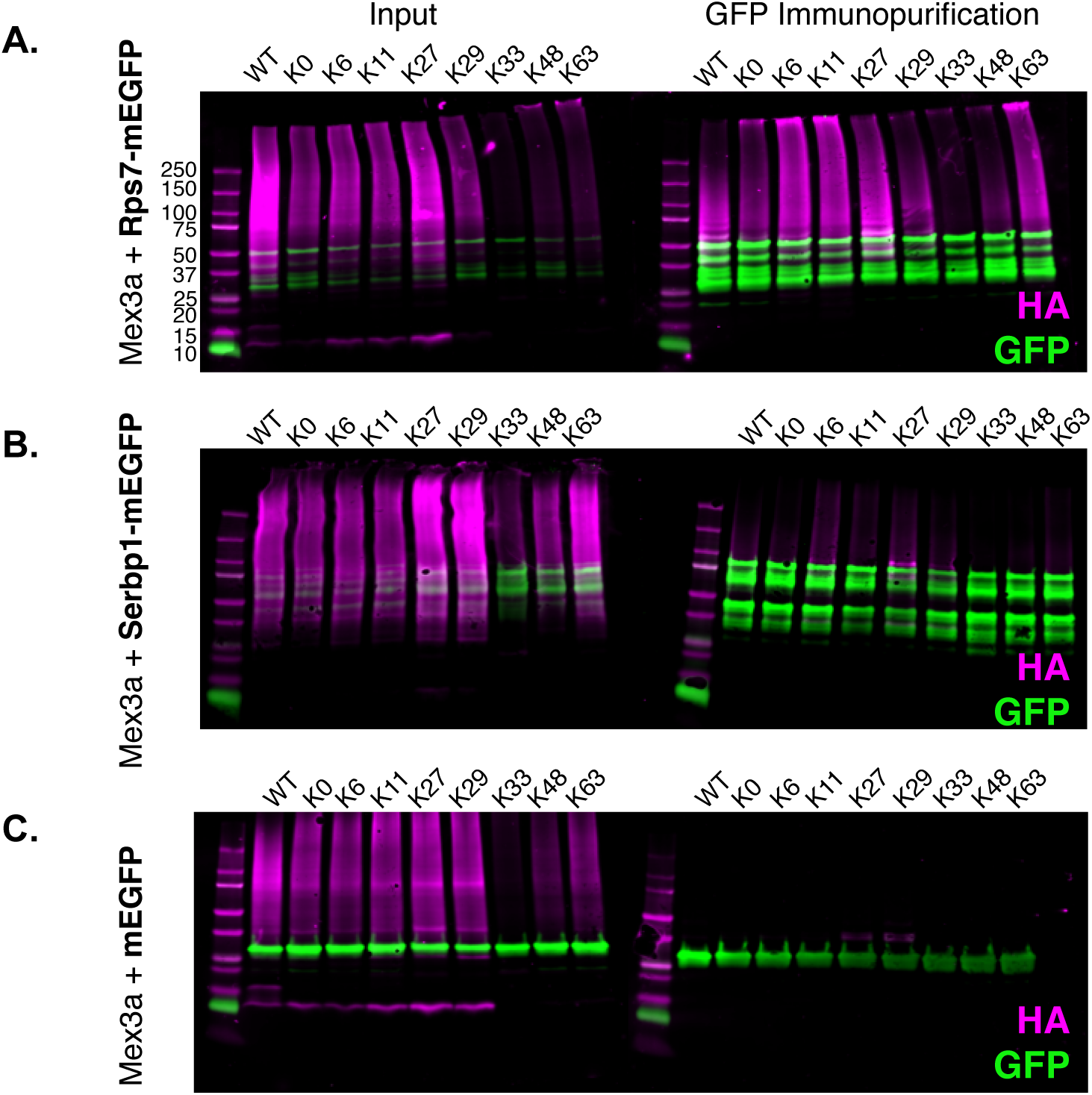
Supporting Main Figure 4. **A.** Control Western blots for results presented in Figure 4E. Left, input lysates taken before immunoprecipitation for transiently transfected HEK293T cells co-transfected with Mex3a and Rps7-mEGFP, HA is shown in magenta and GFP is shown in green. Right, Western blots after immunopurification of GFP constructs, shown in green, and HA-ubiquitin shown in magenta. **B.** same as (A) but for Serbp1-mEGFP. **C.** same as (A) and (B) but for mEGFP construct alone. Due to low expression of K33 ubiquitin construct for all conditions, we do not conclude whether targets are ubiquitinated with K33-linkage or not. All cells were cultured with proteasome inhibitor MG132 to improve K48-ubiquitin abundance.

**Figure S5:**
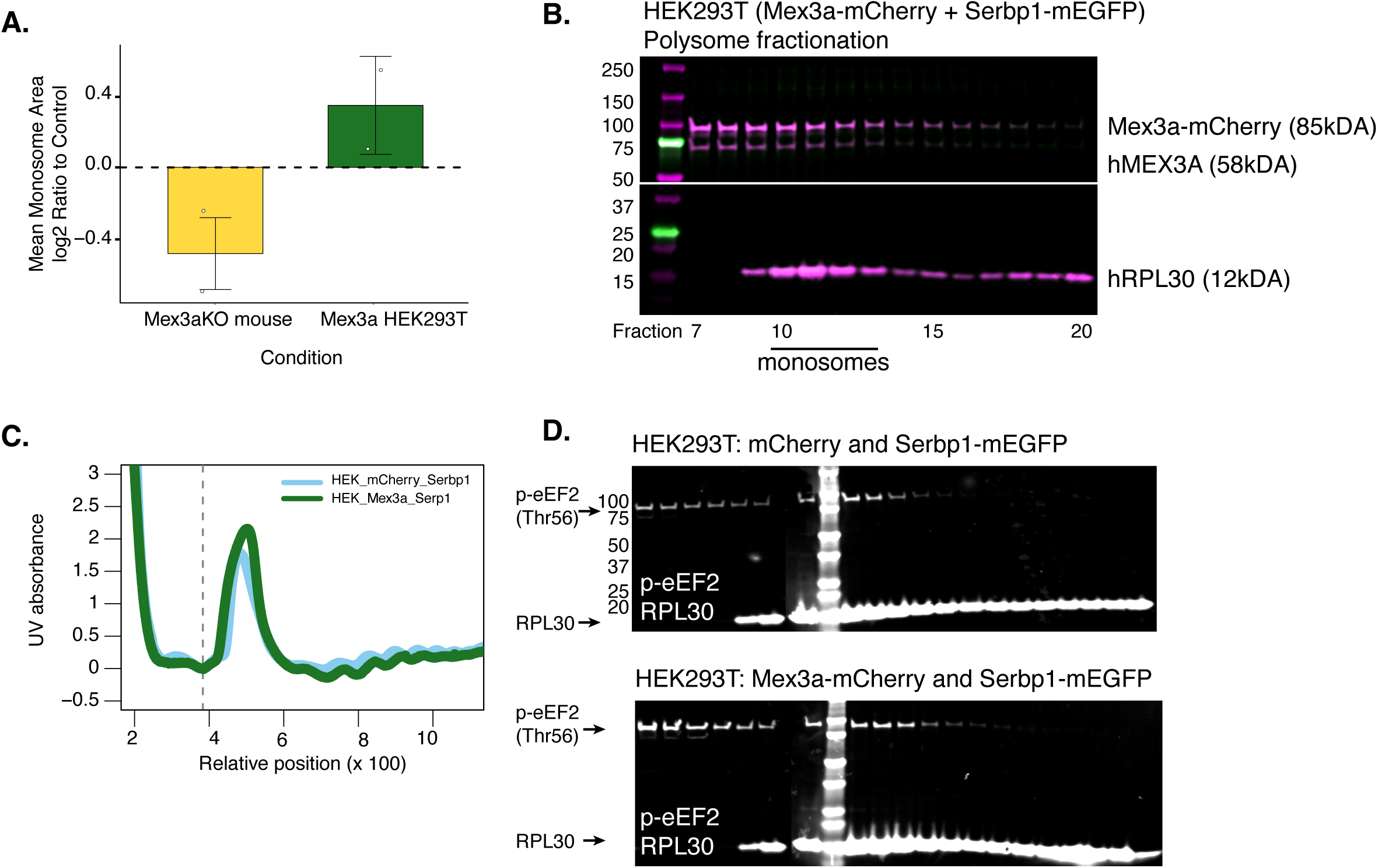
Supporting Main Figure 5. **A.** QuAPPro was used to quantify the area of the monosome peak in each sample, and ratio relative to control was calculated for each experiment. While not significant, there is a trend for decreased monosome area in mouse Mex3a cKO MOE and increased monosome area in HEK293T cells transfected with Mex3a-mCherry **B.** Western blot in HEK293T cells transiently transfected with Mex3a-mCherry and Serbp1-mEGFP. Polysome profiling was performed and a subset of fractions around the monosome was used to blot for Mex3a and RPL30 by Western blot. Westerns for each protein run in parallel with same quantity of sample, images are from different Western blots. **C.** Second replicate of HEK293T polysome profiling experiment, profiles aligned at dotted vertical line and plotted with QuAPPro. **D.** Western blot in polysome fractions around the monosome for endogenous human p-eEF2 in HEK293T cells transfected with mCherry alone and Serbp1-mEGFP (top) or Mex3a-mCherry and Serbp1-mEGFP (bottom). Quantified in Figure 5J.

## Supplemental Files

### Supplemental File 1

Supplemental_File_1_TotalProteome_MassSpec_OMP-GFP_Mex3aWT_cKO.xlsx

*Mass Spectrometry data from total proteomics experiment from OMP GFP+ cells sorted from Mex3a WT and cKO littermates*.

*Raw data available in MassIVE dataset MSV000100330*

### Supplemental File 2

Supplemental_File_2_HyperTRIBE_gg8_results_and_score.xlsx

*Results files and HyperTRIBE scores from gg8-tTA; tetO-Mex3a-HyperTRIBE FACs sorted mCherry+ cells. mCherry negative cells were sorted simultaneously and used as an internal control for each sample*.

*Raw data available in Gene Expression Omnibus (GEO) Accession Number GSE300224*

### Supplemental File 3

Supplemental_File_3_KGG_DIA_Report_pivot.xls

*Mass Spectrometry Spectronaut data of immunopurified proteins after enriching for the Ubiquitin Remnant Motif (K-ε-GG), whole MOE from PN12 littermates, Mex3a WT and cKO*.

*Raw data available in MassIVE dataset MSV000100330*

### Supplemental File 4

Supplemental_File_4_KGG_Spectronaut_Candidates.xlsx

*Spectronaut output comparing ubiquitin post-translational modification levels in Mex3a WT and cKO*.

*Raw data available in MassIVE dataset MSV000100330*

### Supplemental File 5

Supplemental_File_5_DESeq_results_PolysomeRNA_latePoly_vs_earlyPoly.csv

*DESeq results comparing RNA-Seq datasets: late polysome RNA samples vs. early polysome RNA samples for both Mex3a WT and cKO*.

*Raw data available in Gene Expression Omnibus (GEO Accession Number GSE300225*

